# CD4+ T cells regulate sickness-induced anorexia and fat wasting during a chronic parasitic infection

**DOI:** 10.1101/2022.12.17.520896

**Authors:** Samuel E. Redford, Siva Karthik Varanasi, Karina K. Sanchez, Janelle S. Ayres

## Abstract

Catabolic responses of lean and fat energy stores are a component of the host response to infection. Cachexia is an extreme catabolic state characterized by unintentional weight loss and muscle loss, that can include fat loss. Whether cachexia plays any role in host defense or is a maladaptive consequence of host-pathogen interactions remains unknown. Traditionally studies have focused on understanding how inflammatory mediators and cells of the innate immune system contribute to the pathogenesis of cachexia and the depletion of energy stores. The cells of the adaptive immune system that regulate infection-induced cachexia remain elusive. In the present study, we examined the role of the adaptive immune response in cachexia pathogenesis using a murine model of the chronic parasitic infection *Trypanosoma brucei*, the causative agent of sleeping sickness. We found that the cachectic response occurs in two phases with the first stage occurring early in the infection and involved a loss of body fat mass associated with anorexia, and the second stage occurring later in the infection and involved a sustained loss of fat mass that was accompanied by lean mass wasting. CD4+ T cells were necessary for the development of the sickness-induced anorexic response during stage 1 of the infection, which led to adipose triglyceride lipase dependent lipolysis in adipocytes and the resulting fat wasting. Adipose tissue wasting had no impact on host resistance defenses or survival of infection, both of which were antibody-mediated and independent ofCD4+ T cell responses. Our work reveals a new mechanism for infection induced cachexia involving CD4+ T cell regulation of host feeding behavior and an unexpected decoupling of adaptive immune mediated resistance from the cachectic response during infection.

## Introduction

Infections and inflammatory conditions induce a dramatic reprogramming of host metabolic processes. Clinically, the most obvious metabolic response to infection is the wasting of energy stores, however anabolic responses are triggered in addition to catabolic responses. Cachexia is an extreme catabolic state characterized by unintentional weight loss and muscle loss that can include fat loss. Whether cachexia serves a functional role during infections remains unknown but is typically viewed as a maladaptive consequence of host-pathogen interactions. Indeed, in both flies and mice, blocking the wasting of lean and fat tissues protects the host from a diverse array of infections and inflammatory conditions in both invertebrates and vertebrate systems (Clark et al., 2013; Dionne et al., 2006; Schieber et al., 2015). However, mounting a defense response to a pathogen is believed to be energetically costly and requires trade-offs with other biological functions of the host. For example, the acute phase response and antibody production are believed to be as costly as growth and reproductive programs (Abad-Gómez et al., 2013; Amat et al., 2007; Eraud et al., 2005). Therefore, it is conceivable that in the short term, the cachectic response may be beneficial by mediating energy substrate mobilization to meet the energetic costs of the host defense response. However, sustained activation of the cachectic response can become maladaptive contributing to morbidity and mortality of the host.

Cachexia is driven by inflammatory mediators including TNFa, IL-1 and IL-6, which can act directly on energy stores to induce lipolysis and proteolysis of adipose and lean tissues, respectively (Daas et al., 2018a). These cytokines can also mediate metabolic reprogramming during infection by acting on the brain to induce peripheral and behavioral responses including sickness-induced anorexia that control host metabolism. While studies of immune regulation of cachexia have largely focused on the role of cytokines, studies have revealed a role for innate immune cells including macrophages and neutrophils (Petruzzelli et al., 2022; Shukla et al., 2020). In more recent studies, cells of the adaptive immune system, specifically T cells are emerging as a critical regulator of cachexia pathogenesis. In patients with cancer, increased circulating levels of CD8+ T cells and lower levels of regulatory T cells and central memory T cells correlated with increased muscle mass (VanderVeen et al., 2020). In mouse models of cancer cachexia and infections with the parasite *Toxoplasma gondii*, adoptive transfer of CD4+CD44+ T cells or CD4+FOXP3+ Treg cells respectively protected from weight and muscle loss (VanderVeen et al., 2020; Wang et al., 2008) suggesting that CD4+ T cells may play a protective role against both cancer and infection induced cachexia. Whether CD4+ T cells may drive the pathogenesis of cachexia in certain disease contexts remains unknown.

*Trypanosoma brucei* subspecies *brucei* is an extracellular parasitic pathogen that causes African Sleeping Sickness in humans and trypanosomiasis in other mammals. These are chronic infections, that if left untreated, often lead to death of the host. The parasite is vector-borne and transmitted via a tsetse fly bite. The disease is defined by two stages of infection: a blood-born stage where the parasite is circulating at high levels in the blood and a central nervous system (CNS) stage, where the parasite has entered the CNS. The process by which *Trypanosoma brucei* evades the host humoral immune response to establish a chronic infection has been well studied. The parasite is coated with a variant surface glycoprotein (VSG) coat and carries ~2000 copies of the gene but only expresses one copy at a time (Cross et al., 2014). This protein is densely packed onto the surface of the parasite, concealing other cell surface components from immune recognition. Parasites can undergo VSG coat switching, where they begin expressing a different copy of the gene, leading to antigenic variation across the population of parasites in the host (Mugnier et al., 2015). Through this mechanism, they can evade the humoral response due to antibodies not being able to bind to the entire population of the parasite in the host. This leads to the host mounting a continuous humoral response. Furthermore, the parasite has been found to use vascular receptors to localize to adipose tissue and has been reported to induce adipose tissue wasting of the host (De Niz et al., 2021; Trindade et al., 2016). The mechanism by which *T. brucei* induces adipose tissue wasting and whether this is necessary for host defense, or a consequence of the adaptive immune response remains unknown.

In the current study, we utilized a *T. brucei-mouse* infection model to investigate the role of the adaptive immune response in adipose tissue wasting and determine what function this metabolic response has for host defense. We found that the cachectic response occurs in two phases with the first stage occurring early in the infection and involved a loss of body fat mass, and the second stage occurring later in the infection and involved a sustained loss of fat mass that was accompanied by lean mass wasting. The adaptive immune response of the adipose tissue was activated and characterized by a strong Th1 CD4^+^ T cell response, and that correlated with the development of adipose tissue wasting and the sickness induced anorexia response. We found CD4+ T cells to be necessary for the sickness-induced anorexic response that lead to activation of adipose triglyceride lipase (ATGL) in adipocytes, the function of which we found to be necessary for wasting to occur. Using mouse genetic knock outs that lack the major cell types of the adaptive immune system and a mouse model in which *Pnpla2*, the gene encoding ATGL, was knocked out specifically in adipocytes, we revealed a decoupling of host defense from adipose tissue wasting. B cells were necessary for survival of a *T. brucei* infection by promoting host resistance defenses to control parasitemia, while being dispensable for fat wasting. Mice lacking CD4^+^ T cells had no defects in survival or resistance defenses despite being defective in the adipose tissue wasting response. Our study reveals a new role for CD4+ T cells in the development of sickness induced anorexia and adipose tissue wasting and demonstrates that this metabolic response during the acute phase of the infection plays no role in host resistance defenses.

## Results

### *T. brucei* infection activates the adaptive immune response in adipose tissue

*T. brucei* causes a fatal infection in C57BL/6J (WT) mice with a median time to death of 47 days (Figure 1A). Blood parasitemia, increased until ~20 days post-infection where it remained at 10^6^-10^7^ parasites per mL blood for the remainder of the infection (Figure 1B and Supplemental Figure 1A). Consistent with previous reports (Trindade et al., 2016), we found that *T. brucei* infects white adipose tissue (Figure 1C and Supplemental Figure 1B). By 3 days post-infection the gonadal white adipose tissue (GWAT) exhibited ~10^4^ parasites with parasitemia peaking at 4 days post-infection before decreasing towards 6 days post-infection (Figure 1C and Supplemental Figure 1B). Because the levels of adipose tissue parasites were declining, we characterized the immune response to the infection within the GWAT at this time point. We found a significant increase in the percentage of CD45+ cells that were CD4+ and CD8+ T cells in the GWAT of infected mice compared to uninfected controls (Figure 1D). There was a significant increase in the TH1 CD4^+^ T cell populations in the GWAT from infected mice, suggesting that there was a skew towards a more pro-inflammatory immune response (Figure 1E). Consistent with this, the ratio of T regulatory cells (Tregs):TH1 CD4^+^ T cells decreased in infected mice, indicating increased inflammation (Figure 1F). Furthermore, the level of TNFα and IFNγ double positive CD8^+^ T cells was increased during infection (Figure 1G). The level of Ki67 positive CD4 and CD8 T cells were significantly higher during infection, indicating there was increased replication of both T cell populations (Figure 1H). Finally, *T. brucei* infection caused a significant increase in the percentage of CD4 and CD8 T cells in the GWAT that were PD1 and CX3CR1 double positive cells, consistent with increased activation (Figure 1I).

**Figure 1.**
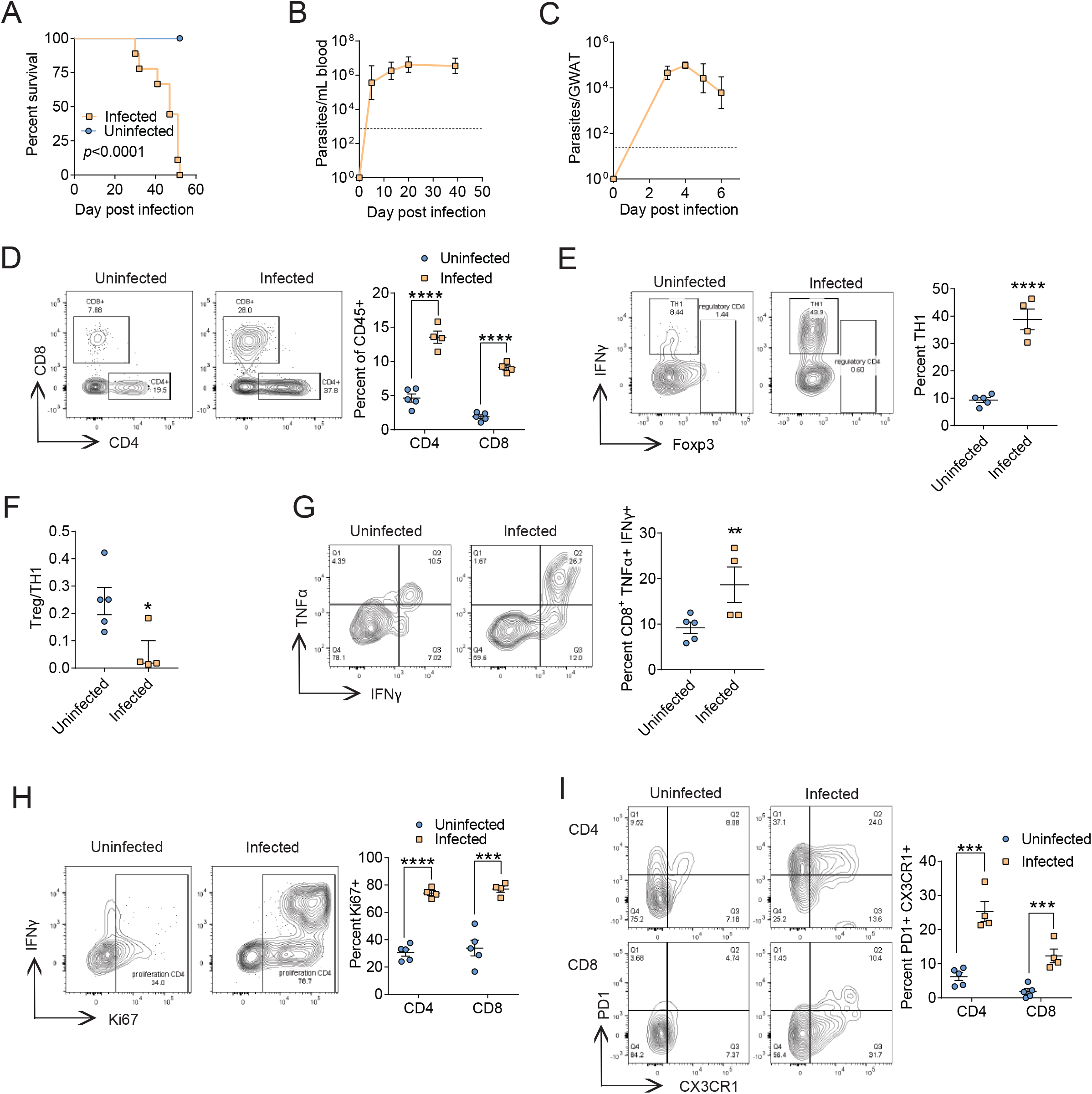
*Trypanosoma brucei* infection activates the adaptive immune response in adipose tissue. 8 week old C57Bl/6 male mice were intravenously infected with 500,000 *T. brucei*. (A) Survival of 8-week-old male C57BL/6J mice infected (n=9) with uninfected age-matched controls (n=9). (B) *T. brucei* levels in blood collected from infected male 8-week-old mice (n=5 for the first two time-point, n=4 for the last two time-points). The limit of detection was 1350 parasites per mL of blood and is represented by the dotted line on the graph. (C) *T. brucei* levels in gonadal white adipose tissue (GWAT) from infected C57BL/6J mice (n=5 for days 3 and 5, n=4 for days 4 and 6). The limit of detection was 3400 parasites per gram of tissue, represented by the dotted line on the graph. (D) Percentage of CD45^+^ CD3^+^ cells that were either CD4^+^ or CD8^+^ T cells from single cells isolated from GWAT of either infected (n=4) or uninfected C57BL/6J mice (n=5) at 6 days post-infection. (E) Percentage of CD4^+^ T cells shown in panel D that were IFNγ^+^. (F) Percentage of CD3^+^ CD4^+^ T cells from panel D that were Foxp3^+^ divided by TH1 T cells in panel E for a ratio of inflammation. (G) Percentage of CD8^+^ T cells from panel D that were TNFα^+^ and IFNγ^+^. (H) Percentage of CD4^+^ and CD8^+^ T cells from panel D that were Ki67^+^. (I) Percentage of CD4^+^ and CD8^+^ T cells from panel D that were PD1^+^ and CX3CR1^+^. * p=0.0635, ** p<0.05, *** p<0.01, and *** p<0.0001 All statistical tests for pairwise comparisons were unpaired Student’s T-test or Mann Whitney test between uninfected and infected of the same condition. For survival, Log-rank analysis. Error bars represent +/- SEM except for (B) and (C) which is plotted as geometric mean with error bars showing geometric SD. Frequencies displayed on gates are percent of parent gate which may be different than what is being displayed on graph which is listed above for each panel. Data in A are two independent experiments combined and data in B-I represent one independent experiment.

We next profiled the proportion of natural killer (NK) cells and found that while infection caused a decrease in the level of NK cells in the GWAT, they produced more IFNγ (Supplemental Figure 1CD). We found no difference in the proportion of GWAT macrophages between infected and uninfected mice (Supplemental Figure 1E). GWAT macrophages from infected mice had higher expression of MHCII but lower median fluorescent intensity of TNFα compared to uninfected mice, suggesting the increased inflammation we observed was not universal and instead was specific to T cells (Supplemental Figure 1F and 1G). Taken together, these results indicate that *T. brucei* infection causes an increase in the levels of activated CD4+ and CD8+ T cells in the adipose tissue that correlates with the decline in fat parasite burdens.

### *T. brucei* infection causes stage specific depletion of fat and lean energy stores

We analyzed the weight loss kinetics of *T. brucei* infected mice and found that they lost ~15% body weight by day 5-6 post-infection (Figure 2A and Supplemental Figure 2A). By ~12 days post-infection, infected mice regained their weight to levels comparable to uninfected control animals and then again exhibited weight loss by ~20 days post-infection that was sustained for the remainder of the infection (Figure 2A and Supplemental Figure 2A). We performed MRIs on mice over the course of the infection and found that the weight loss observed at the early stage of infection between days 5 and 6 post-infection was due to a loss of fat mass (Figure 2B and Supplemental Figure 2B). Infected mice did not exhibit a loss of lean mass until 20-days post-infection (Supplemental Figure 2C-D). In agreement with our MRI analysis, we found that between 5- to 6-days post-infection, infected mice had significantly less GWAT and IWAT compared to uninfected control mice (Figure 2C-D and Supplemental Figure 2E-F). This reduced fat mass persisted for the duration of the infection (Figure 2C-D and Supplemental Figure 2E-F). Measurements of the gastrocnemius leg muscle revealed, similar to our MRI analysis, that lean mass was similar between infected and uninfected at mice at day 5-6 post-infection, and loss of lean mass occurred at later stages of infection (Supplementary Figure 2G-H). Taken together, *T. brucei* induced cachexia involves two distinct stages with the first stage occurring early in the infection and involving a loss of body fat mass. Stage 2 occurs later in the infection and involves a sustained loss of fat mass that is accompanied by lean mass wasting.

**Figure 2.**
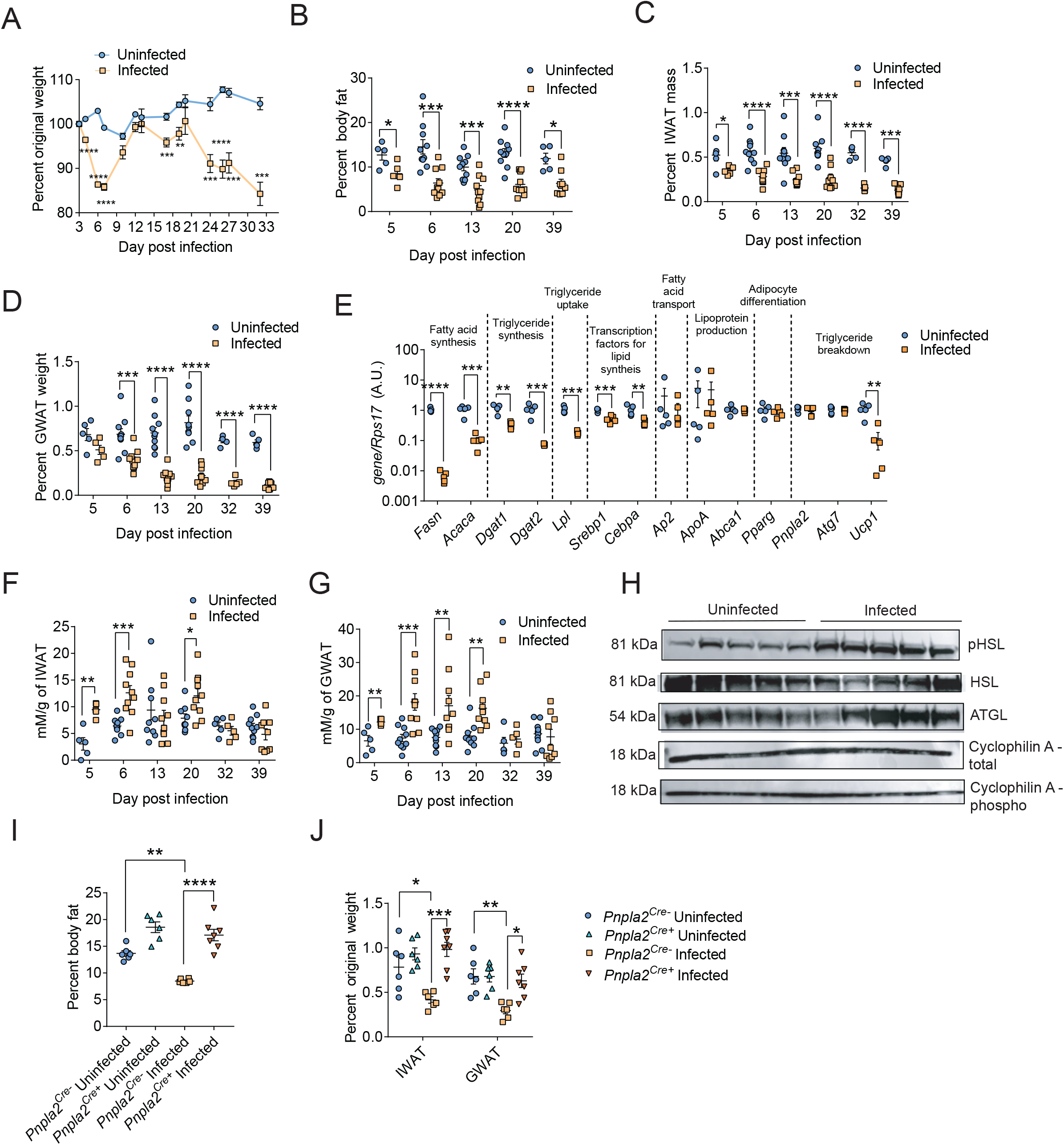
Mice infected with *Trypanosoma brucei* undergo lipolysis-dependent adipose tissue wasting. 8 week old C57Bl/6 male or female mice were intravenously infected with 500,000 *T. brucei*. (A) Percent of original weight was determined. The weight just prior to the time of infection was used as the original weight. Infected (n=25) and uninfected mice (n=25). Mice were used for dissections at timepoints shown in next panel and so n was reduced by 5 at each of those timepoints. In addition, due to time constraints, on days with a dissection, only the mice used for the dissection were weighed. (B) Percent total body fat of uninfected (n=5 for 5 days post-infection, n=10 for each other timepoint) or infected (n=5 for 5 days post-infection, n=10 for each other timepoint) mice as a percentage of their pre-study weight. (C) Inguinal white adipose tissue (IWAT) weights from same mice as in panel B normalized to the pre-study weight of each mouse. (D) GWAT weights from mice in panel B normalized to the pre-study weight of each mouse. (E) Gene expression of shown genes in IWATs from uninfected (n=5) and infected (n=5) C57BL/6J mice normalized to *Rps17* at 6 days postinfection. (F) Amount of free fatty acids (FFAs) released from IWAT from uninfected (n=5) or infected (n=5) mice incubated in Krebs-Ringer bicarbonate Hepes (KRBH) buffer normalized to the mass of IWAT put in the buffer. (G) Amount of FFAs released from GWAT isolated from uninfected (n=5) or infected (n=5) mice and incubated in KRBH buffer, normalized to the GWAT weight placed in the buffer. (H) Western blot of hormone-sensitive lipase (HSL), phospho-HSL, ATGL, and Cyclophilin A of infected mice (n=5) or uninfected mice (n=5). Blot labeled “Cyclophilin A total” is the loading control for HSL and ATGL blots. Blot labeled “Cyclophilin A phospho” is the loading control for pHSL blot. Protein size is listed next to each protein. (I) Percent body fat of *Pnpla2^flox/flox^;Fabp4^Cre+^* (*Pnpla2^Cre+^*) mice that were infected (n=7) or uninfected (n=6) compared to *Pnpla2^flox/flox^;Fabp4^Cre-^* (*Pnpla2^Cre-^*) littermates that were infected (n=6) or uninfected (n=6) normalized to pre-study weight of each mouse at 6 days post-infection. (J) Same mice as in H showing both GWAT and IWAT fat pad weights normalized to the pre-study weight of each mouse. * p<0.05, ** p<0.01, *** p<0.001, and *** p<0.0001. For pairwise comparisons, Unpaired Student’s T-test, Mann-Whitney Test or One-Way ANOVA with post Tukey’s test. Error bars represent +/- SEM. Panels A, H, and I show one independent representative experiment while the other panels show two independent experiments combined.

Adipose tissue stores are regulated by lipogenesis and lipolysis pathways. We characterized how *T. brucei* infection influenced lipogenesis and lipolysis of the WAT. At 6 days post-infection, WAT from *T. brucei* infected mice had significantly reduced expression of genes involved in lipogenesis that was sustained during the infection (Figure 2E and Supplemental Figure 2I). Using an ex vivo lipolysis assay, we found that both IWAT and GWAT fat deposits isolated from *T. brucei* infected mice at days 5-, 6-, 13-, and 20-days post-infection released significantly more amounts of FFA compared to uninfected mice, with the highest levels observed at day 6 post-infection (Figure 2F and 2G). Lipolysis is regulated by adipose triglyceride lipase (ATGL), which is encoded by the gene *Pnpla2* and is the first enzyme and rate limiting step of the pathway, as well as hormone-sensitive lipase (HSL). *T. brucei* infection did not cause a change in the levels of *Pnpla2* mRNA, however the majority of mice examined had increased ATGL protein compared to uninfected mice in the adipose tissue. *T. brucei* infection also caused an increase in the levels of activated HSL (phospho-HSL) in the adipose tissue compared to uninfected mice at day 6 post-infection (Figure 2H and Supplemental Figure 2J). To test the importance of lipolysis for *T. brucei*-induced adipose tissue wasting, we infected *Pnpla2^flox/flox^;Fabp4^cre+^* (*Pnpla2^Cre+^*) in which the gene encoding ATGL, *Pnpla2*, has been deleted specifically from adipocytes and compared their fat wasting phenotype to infected *Pnpla2^flox/flox^;Fabp4^cre-^* (*Pnpla2^Cre-^*) littermates. We focused our analysis specifically on day 6 postinfection as this is the earliest time point, we observed fat wasting and that correlates with declines in WAT parasitemia. Infected *Pnpla2^Cre+^* mice were protected from adipose tissue wasting whereas infected *Pnpnla2^Cre-^* littermates lost around 40 percent of their fat mass as indicated by MRI (Figure 2I and Supplemental 2K). Furthermore, *Pnpnla2^Cre+^* infected mice were protected against wasting of their GWAT and IWAT deposits (Figure 2J). These results indicate that the adipose tissue wasting induced by a *T. brucei* infection is due to changes in both the lipogenic and lipolytic programs of the WAT, and that ATGL is necessary for infection-induced adipose tissue wasting that occurs in the early stage of infection.

### Adaptive immune activation in adipose tissue is independent of wasting and lipolysis

Because adaptive immune activation correlated with adipose tissue wasting in the early stage of infection, we asked if and how these processes were related during infection. First, we asked if lipolysis and adipose tissue wasting were necessary for immune activation. We infected *Pnpnla2^Cre+^* and *Pnpnla2^Cre-^* littermates and isolated the GWATs at day 6 post-infection to profile immune cells as we did in Figure 1. Infected *Pnpnla2^Cre+^* mice had significantly more CD45^+^ cells in the GWAT than the infected *Pnpnla2^Cre-^* mice (Supplemental Fig. 3A). However, *Pnpnla2^Cre+^* mice had significantly more adipose tissue during infection (Figure 2I). Upon normalization of cell numbers to the mean of GWAT mass for each group, we found no significant difference between infected *Pnpnla2^Cre+^* or infected *Pnpnla2^Cre-^* mice. This indicates that wasting does not change the number of lymphocytes present when controlling for the mass of adipose tissue (Figure 3A). The proportion of CD45^+^ cells that were either CD4^+^ and CD8^+^ T cells increased to comparable levels in both *Pnpnla2^Cre+^* and *Pnpnla2^Cre-^* infected mice with the strongest increase in CD4+ T cells, mimicking our earlier results in C57Bl/6 mice (Figure 1D and 3B). We found a comparable increase in IFNγ in CD4^+^ T cells and a comparable decrease in the ratio of Tregs:Th1 CD4^+^ T cells from both infected *Pnpnla2^Cre+^* and infected *Pnpnla2^Cre-^* mice compared to uninfected controls (Figure 3C-D) without an increase in IL17, or IL4 (Supplemental Figure 3B-C) indicating *T. brucei* triggers a predominantly Th1 response in the adipose tissue independent of lipolysis and fat wasting. Furthermore, there was no difference in the levels of IFNγ and TNFα double positive CD8+ T cells or Ki67 staining in both CD4^+^ and CD8^+^ T cells between infected *Pnpnla2^Cre+^* and infected *Pnpnla2^Cre-^* mice (Figure 3E and Supplemental Figure 3D). We also found no difference in the proportion of CD45+ cells that were B cells or in IgD or MHCII expression on the B cells indicating no strong changes in B cell levels or activation (Supplemental Figure 3E-F). *T. brucei* infection induced a comparable level of IFNγ production by NK cells (Supplemental Figure 4A-B). Finally, from our characterization of myeloid cells, we saw the levels of macrophages in the GWAT trended lower in infected *Pnpnla2^Cre-^* mice compared to uninfected controls. In *Pnpnla2^Cre+^*, uninfected mice exhibited lower levels and macrophage levels did not significantly change upon infection (Supplemental Figure 4C). Macrophage expression of TNFα in the GWATs from both infected *Pnpnla2^Cre+^* and *Pnpnla2^Cre-^* mice decreased to comparable levels compared to uninfected controls (Supplemental Figure 4D). We also saw no differences in macrophage polarization caused by either the infection or adipose tissue wasting (Supplemental Figure 4E). Taken together, adipose tissue lipolysis and wasting are not necessary for *T. brucei* activation of the adaptive immune response in adipose tissue.

**Figure 3.**
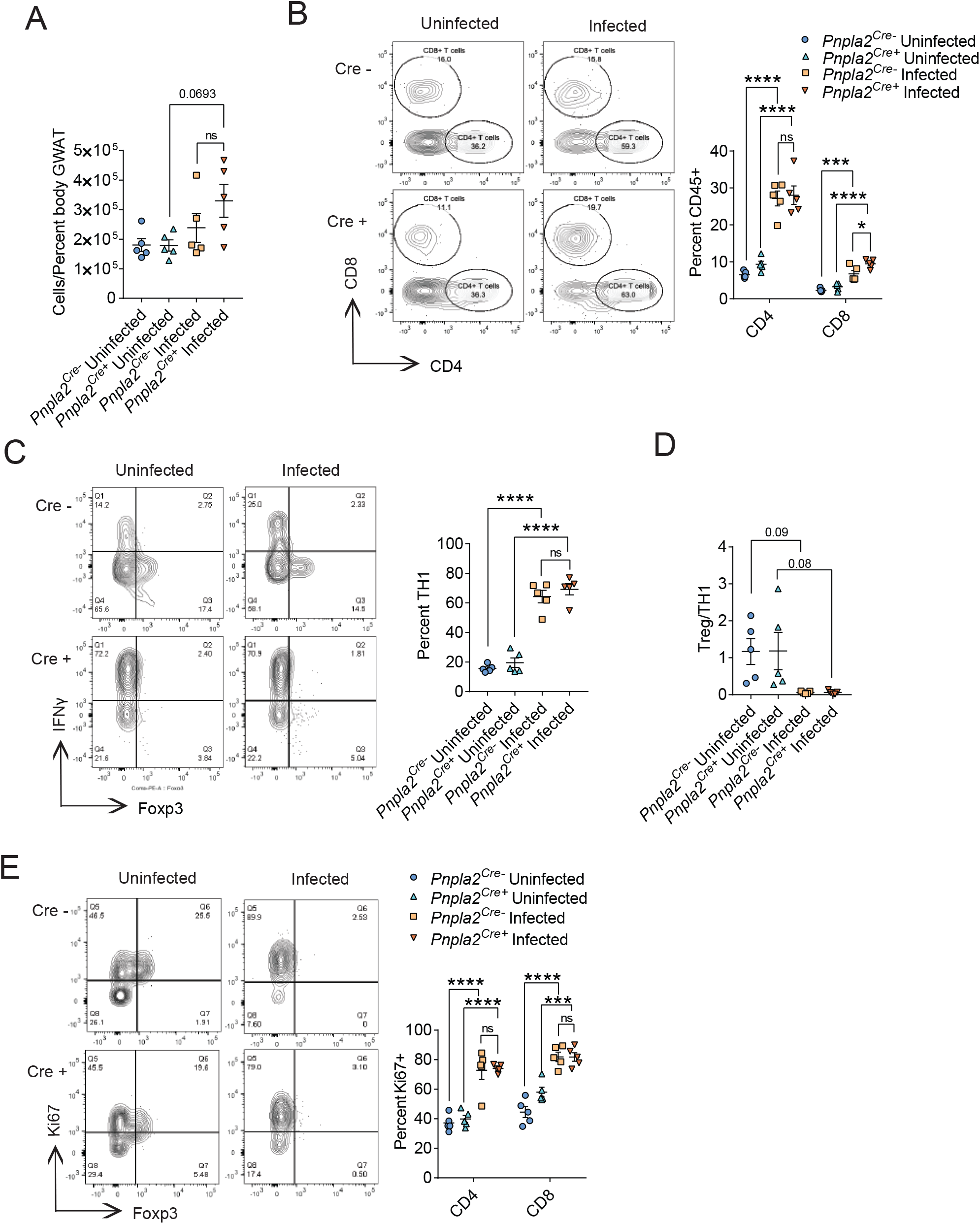
Adipose tissue wasting is not necessary for activation of adaptive immunity in adipose tissue. 8 week old female mice were intravenously infected with 500,000 *T. brucei* and the immune cells profiled. Cells were isolated from gonadal white adipose tissue (GWAT) of uninfected *Pnpla2^flox/flox^;Fabp4^Cre+^* (*Pnpla2^Cre+^*) mice (n=5), uninfected *Pnpla2^flox/flox^;Fabp4^Cre-^* (*Pnpla2^Cre-^*) (n=5), infected *Pnpla2^Cre-^* (n=5) and infected *Pnpla2^Cre+^* (n=5) littermates at 6 days post-infection and immune cell populations were quantified (A) The total number of CD45^+^ cells isolated from fat pads normalized to average percent GWAT mass for each group. (B) Percent of CD45^+^ cells that are CD4^+^ or CD8^+^ T cells. (C) Percent of CD3^+^ CD4^+^ positive for IFNγ. (D) Ratio of the percent of CD4^+^ T cells that were Foxp3^+^ to the percent of CD4^+^ T cells that were Foxp3^-^ and IFNγ^+^. (E) Percent of either CD4^+^ or CD8^+^ T cells that were positive for Ki67. * p<0.05, ** p<0.01, *** p<0.001, and *** p<0.0001 One-way ANOVA comparing mean to every other mean with Tukey correction for multiple comparisons. Error bars represent +/- SEM. Flow charts are presented to show gating with Uninfected *Pnpla2^Cre-^* mouse upper-left, uninfected *Pnpla2^Cre+^* mouse lower-left, infected *Pnpla2^Cre-^* mouse upper-right, and infected *Pnpla2^Cre+^* mouse lower-right. Frequencies displayed on gates are percent of parent gate which may be different than what is being displayed on graph which is listed above for each panel. One representative experiment shown for each panel.

We next determined if lipolysis and adipose tissue wasting was necessary for the immune response outside of the adipose tissue during a *T. brucei* infection. Our immune phenotyping revealed differences in the cell proportions within the spleen compared to adipose tissue that were independent of lipolysis and adipose tissue wasting. Spleens from both infected *Pnpnla2^Cre+^* and *Pnpnla2^Cre-^* mice did not exhibit an increase in the total cell numbers at day 6 post-infection (Supplemental Figure 5A), however we did observe a drop in both CD4+ and CD8+ T cells (Supplemental Figure 5B). IFNγ+ CD4+ and CD8+ T cells increased in the spleens from both infected *Pnpnla2^Cre+^* and *Pnpnla2^Cre-^* mice (Supplemental Figure 5C). While splenic CD4+ T cells exhibited an increase in ki67 during infection, CD8+ T cells exhibited a decrease in ki67 levels (Supplemental Figure 5D). Finally, we observed that the number of splenic B cells increased, while NK cells decreased (Supplemental Figure 5E and 6A). Splenic macrophages also increased with a drop in TNFα production and an increase in MHCII expression (Supplemental Fig. 6B-D). Taken together, our data indicate that *T. brucei* infection causes distinct immune cell profiles and activation in the spleen and adipose tissue in a lipolysis and fat wasting independent manner.

### CD4^+^ T cells are necessary for adipose tissue wasting

We next asked if the adaptive immune response was necessary for *T. brucei* induced adipose tissue wasting. Rag1 KO mice, which lack mature T and B cells, were highly susceptible to a *T. brucei* challenge with 100% mice succumbing to the infection by ~20 days post-infection and exhibiting an order of magnitude higher blood parasitemia compared to infected wild type mice (Figure 4A-B). Despite this increased susceptibility, we found from our MRI analysis that the severity of adipose tissue wasting was significantly less in infected Rag1 KO mice compared to infected wild type mice (Figure 4C and Supplemental Figure 7A). Furthermore, from direct measurements of fat deposits we found that infected Rag1 KO mice were completely protected from wasting of their GWAT and IWAT deposits while wild type infected mice lost ~50% or more of their IWAT and GWAT mass respectively compared to uninfected wild type controls (Figure 4D and Supplemental Figure 7B). These results indicate that the adaptive immune system is necessary for *T. brucei* induced adipose tissue wasting to occur.

**Figure 4.**
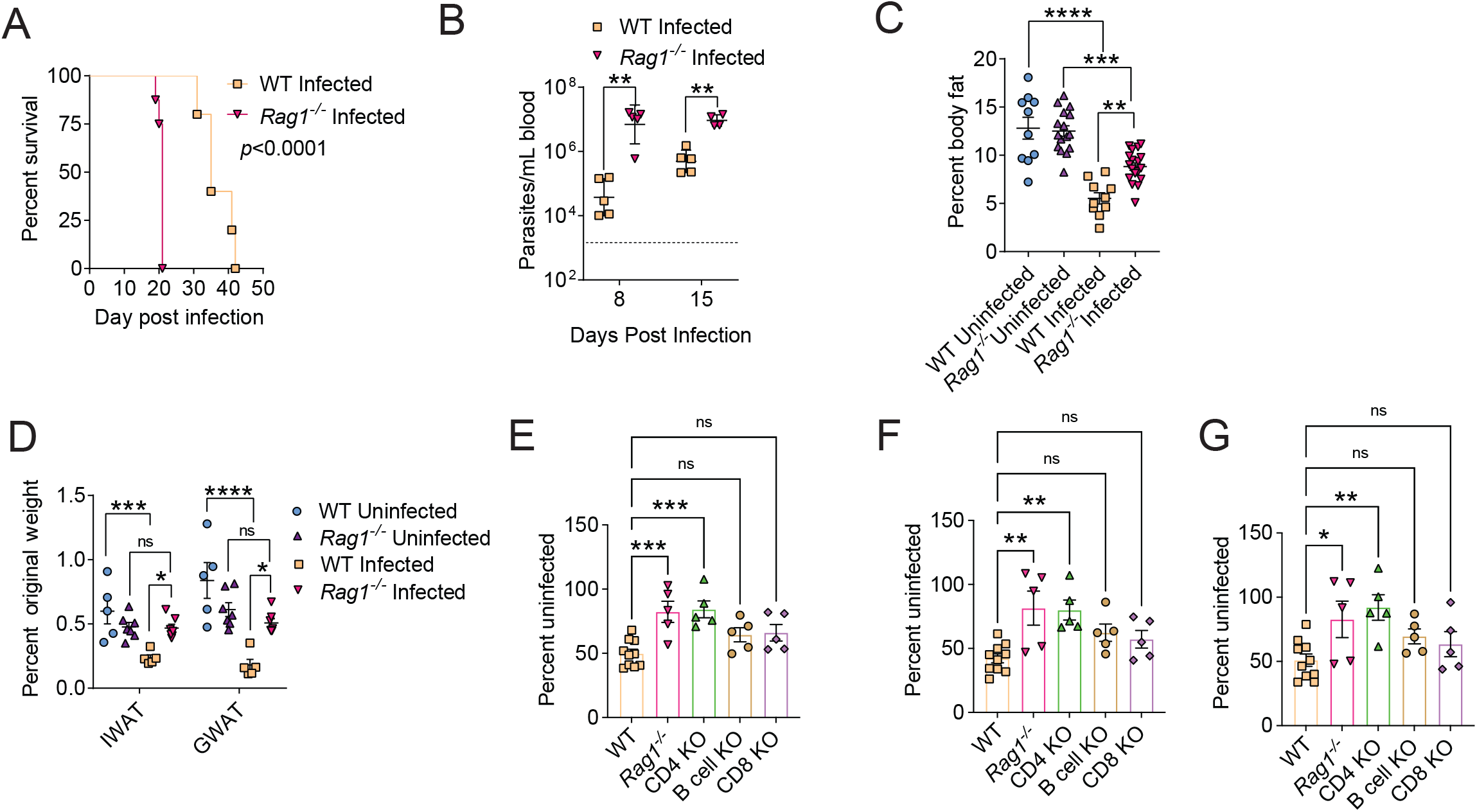
CD4+ T cells are necessary for *T. brucei* induced adipose tissue wasting. (A) Survival of male 8-week-old B6.129S7-Rag1^tm1Mom^/J mice (*Rag1*^-/-^) (n=8) compared to male age-matched wild type controls (WT) (n=5). (B) Blood parasitemia of infected *Rag1*^-/-^ (n=5 at 8 days post-infection, n=4 at 15 days post-infection) and WT (n=5 for each time-point). The limit of detection was 1350 parasites/mL and is displayed by the dotted line. (C) Percent body fat of uninfected (n=10) and infected (n=10) WT and uninfected (n=15) and infected (n=18) *Rag1*^-/-^ mice. (D) Percent IWAT and GWAT weights from uninfected (n=5) and infected (n=5) WT mice compared to uninfected (n=7) and infected (n=7) *Rag1*^-/-^mice normalized to mouse weight at time 0hr. (E-G) Body fat characterization of Infected (n=10) WT mice compared to infected B6.129S7-Rag1^tm1Mom^/J mice (*Rag1*^-/-^) (n=5), infected B6.129S2-Cd4^tm1Mak^/J (CD4 KO) (n=5), infected B6.129S2-Cd8a^tm1Mak^/J (CD8 KO) (n=5), and infected B6.129S2-Ighm^tm1Cgn^/J (B cell KO) (n=5). (E) MRI of fat mass was done, and the infected mice were normalized to the mean of the fat mass of the uninfected mice for each genotype. (F) Same mice as in E were dissected and inguinal white adipose tissue (IWAT) was weighed to confirm MRI results. Displayed is the weight of the infected mice normalized to the mean of the uninfected mice for each genotype. (G) Same mice as in E showing gonadal white adipose tissue (GWAT) of the infected mice normalized to the mean of the uninfected for each genotype. All infections were carried out with an intravenous injection of 500,000 parasites. * p<0.05, ** p<0.01, *** p<0.001, and *** p<0.0001 For A, the survival curve was analyzed with Log-rank (Mantel-Cox) test. For panel B, unpaired Student T-tests were performed between *Rag1* KO mice and WT at each time point. For the remainder of the panels, One-way ANOVA were with a post Tukey correction for multiple comparisons. Error bars represent +/- SEM except in panel B which has geometric mean plotted and geometric SD as the error bars. Panel D shows two independent experiments combined, all other panels show one representative experiment.

To determine which cell type of the adaptive immune system was necessary for the adipose tissue wasting in the early stage of infection, we employed genetic KO mice that lack each of the major cell types within the adaptive immune system and characterized their adipose tissue wasting response during a *T. brucei* challenge at day 6 post-infection. Under uninfected conditions, there are trends towards baseline differences in the adiposity of mice that lack different cellular components of the adaptive immune system (Supplemental Figure 7C). To account for this, we normalized the weights of infected mice to that of uninfected mice of the same genotype that were weight matched at the time of infection (Supplemental Figure 7D). There was no significant difference in the amount of fat loss between infected mice deficient for B cells, CD8+ T cells and wild type mice (Figure 4E-G and Supplemental Figure 7E-G). By contrast, we found that infected CD4^+^ T cell KO mice had significantly less body fat loss compared to infected wild type mice and were protected from adipose tissue wasting to the same degree as the *Rag1* KO mice (Figure 4E-G and Supplemental Figure 7E-G). Taken together, our data suggest that CD4+ T cells are the cell type responsible for driving adipose tissue wasting during *T. brucei* infection.

### CD4^+^ T cell dependent sickness-induced anorexia drives adipose tissue wasting

Many pathogens cause a sickness-induced anorexic response in their host that results in a reduced consumption of food during the infection. Wild type mice infected with *T. brucei* exhibit a robust anorexic response early in the infection that begins at ~4 days post-infection, peaks at 6 days post-infection and returns to feeding levels of uninfected mice by ~8-10 days post-infection (Figure 5A). Because this anorexic response coincides with the fat wasting phenotype, we observe early in the infection course of *T. brucei* infected mice, we hypothesized that the mechanism by which CD4+ T cells mediate adipose tissue wasting is through the induction of the sickness-induced anorexic response. We found that both Rag1 KO and mice deficient for CD4+ T cells were significantly protected from developing the sickness-induced anorexic response, while infected mice deficient for B and CD8+ T cells exhibited comparable severity and kinetics of the anorexic response as infected wild-type mice (Figure 5B-C and Supplemental Figure 8A). Thus CD4+ T cells are necessary to drive the sickness-induced anorexic response during *T. brucei* infection.

**Figure 5.**
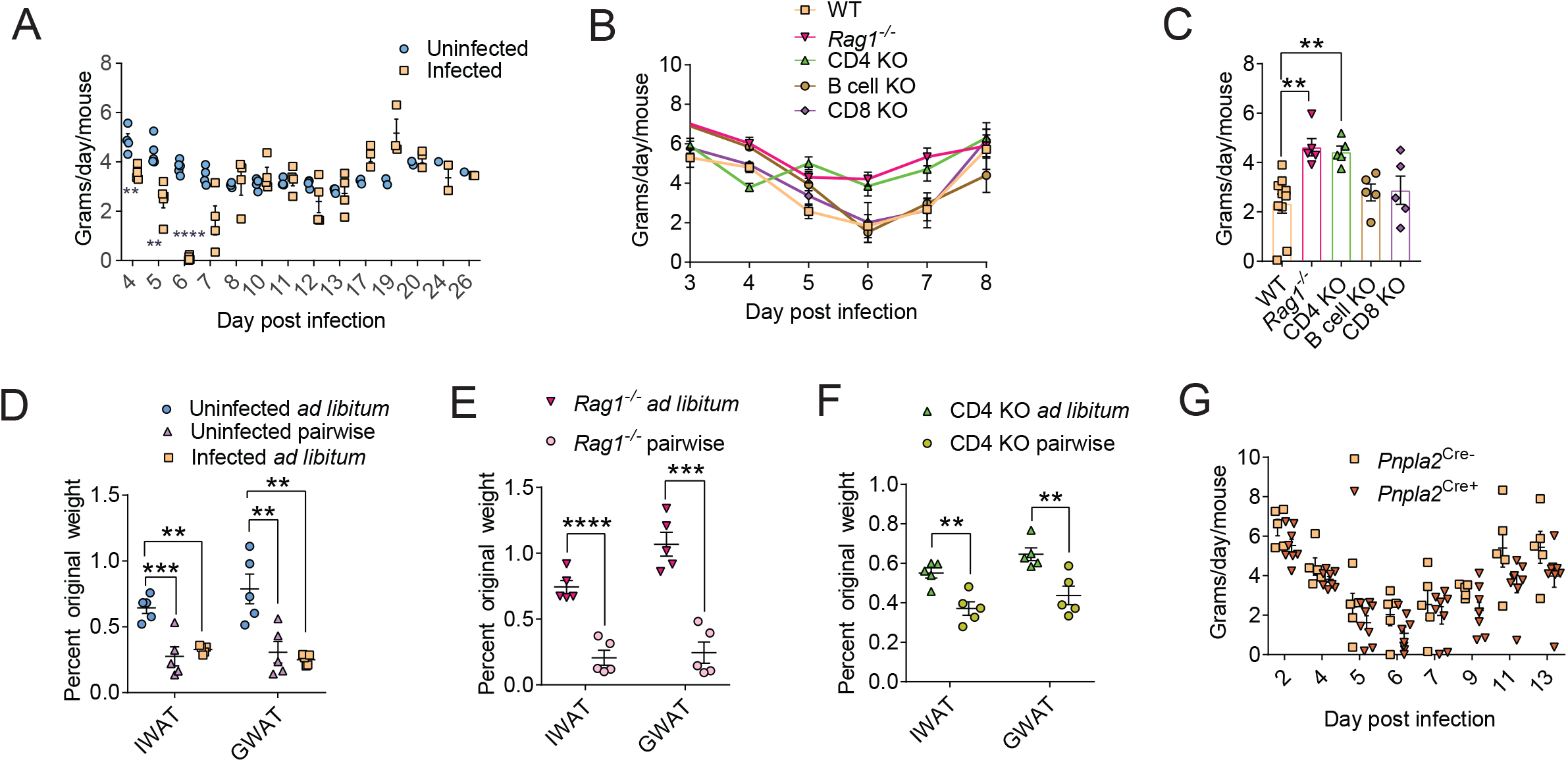
CD4^+^ T cells are necessary for the sickness induced anorexic response. (A) 8 week old C57BL/6J mice were infected with 500,000 parasites and food consumption was measured and compared to uninfected control animals. Mice were group housed and the average food consumed per mouse was determined by dividing the overall food intake of the cage by the number of mice in the cage for each day. Infected (n=5-25 mice, 1-5 cages depending on time point) and uninfected (n=10-25 mice, 2-5 cages depending on time point). (B) Food intake of single housed infected WT (n=5-10) mice, B6.129S7-Rag1^tm1Mom^/J (n=5) mice (*Rag1*^-/-^), B6.129S2-Cd4^tm1Mak^/J (n=5) mice (CD4 KO), B6.129S2-Cd8a^tm1Mak^/J (n=5) mice (CD8 KO), and B6.129S2-Ighm^tm1Cgn^/J (n=5) mice (B cell KO). (C) Average food intake of mice in panel B over days 4-7 post-infection. (D) Weights of both inguinal white adipose tissue (IWAT) and gonadal white adipose tissue (GWAT) from single caged, uninfected C57BL/6J fed ad libitum (n=5), uninfected C57BL/6J mice pairwise fed and given only the amount of food infected mice eat (n=5), and infected C57BL/6J fed ad libitum (n=5) dissected at 7 days post-infection. (E) IWAT and GWAT weights normalized to starting body weight of 12 week old *Rag*^-/-^ mice infected and fed *ad libitum* (n=5) or infected and pairwise fed to match the amount consumed by WT infected mice as shown in Figure 5A (n=5). (F) IWAT and GWAT weights normalized to starting body weight of 8 week old CD4 KO mice infected and fed *ad libitum* (n=5) or infected and pairwise fed to match the amount consumed by WT infected mice as shown in Figure 5A (n=5). (G) Food intake of single housed infected *Pnpla2^flox/flox^;Fabp4^Cre+^* (*Pnpla2^Cre+^*) mice (n=7) and infected *Pnpla2^flox/flox^;Fabp4^Cre-^* (*Pnpla2^Cre-^*) littermates (n=5). * p<0.05, ** p<0.01, *** p<0.001, and *** p<0.0001 unpaired Student’s T-test for panels A, E, F for each time point or fat pad. For other panels, One-way ANOVA comparing the mean of each column to every other mean using Tukey correction for multiple comparisons was used. Error bars represent +/- SEM. One representative experiment shown for each panel.

Our data thus far demonstrate that CD4+ T cells drive both sickness-induced anorexia and adipose tissue wasting in *T. brucei* infected hosts. We first attempted to test the importance of this anorexic response for adipose tissue wasting using a force feeding regimen (Rao et al., 2017). However, we found that when we force fed wild type mice infected with *T. brucei*, their anorexic response became even more severe, and they consumed even less chow (data not shown). As an alternative approach, to determine if the reduced food consumption due to the anorexic response was sufficient to cause adipose tissue wasting, we performed a pairwise feeding experiment where we fed uninfected wild type mice the same amount of food that the infected wild type mice eat, mimicking the anorexic response in the infection. We found that the uninfected pairwise fed mice exhibited the same extent of adipose tissue wasting as infected mice fed *ad libitum* as indicated by no differences in IWAT and GWAT weights at day 7 post-infection (Figure 5D and Supplemental Figure 8B). Furthermore, when we fed Rag1 KO mice and CD4+ T cell deficient mice the same amount of food as infected wild type mice early in the infection we found that both *RAG1* KO mice and CD4^+^ T cell KO mice lost a significant amount of fat compared to infected mice fed *ad libitum*, indicating that reduced food intake is sufficient to induce adipose tissue wasting in Rag1 KO and CD4 T cell deficient infected mice (Figure 5E-F and Supplemental Figure 8C-D). To confirm that anorexia acts upstream of lipolysis and adipose tissue wasting, we infected *Pnpla2 Cre+* and *Cre-* littermates with *T. brucei* to determine how lipolysis and adipose tissue wasting influence the anorexic response. We found that *Pnpla2 Cre+* mice developed an anorexic response that was comparable in kinetics and severity to infected *Pnpla2 Cre-* littermates that are able to undergo lipolysis, indicating that sickness-induced anorexia is not downstream of lipolysis and adipose tissue wasting (Figure 5G). Taken together, CD4+ T cells are a driver of sickness-induced anorexia during *T. brucei* infection that causes lipolysis and adipose tissue wasting.

### Decoupling adaptive immune system mediated resistance from the metabolic response to infection

Whether adipose tissue wasting plays a functional role in host defense during infection remains unclear. Previous work showed that the humoral response is critical in controlling *T. brucei* infections (Ponte-Sucre, 2016). To formally test whether lipolysis and adipose tissue wasting is important for host defense, we first quantified the amount of *T. brucei* specific antibodies circulating in our genetic mouse models. Both Rag1 KOs and mice deficient for B cells failed to produce *T. brucei* specific IgG or IgM (Figure 6A and Supplemental Figure 9A-B). By contrast, mice that were protected from undergoing lipolysis and adipose tissue wasting including *Pnpla2 Cre+* and CD4 KO mice were able to mount an antibody response when challenged with *T. brucei* (Figure 6A-B and Supplemental Figure 9A-B). Mice deficient for CD8 T cells were also capable of generating IgG and IgM that bound to *T. brucei* comparable to what we observed in infected wild type mice (Figure 6A and Supplemental Figure 9A-B). In agreement with our antibody data, Rag1 KO and B cell KO mice exhibited higher blood and adipose tissue parasitemia compared to wild type mice, suggesting they had a compromised resistance response to the parasite (Figure 6C-D). Infected CD4 KO and *Pnpla2 Cre+* mice exhibited no differences in blood or adipose tissue parasitemia compared to their respective controls demonstrating that adipose tissue wasting is not necessary for the host resistance response to the parasite systemically or locally in the fat (Figure 6C-F). CD8 KO mice also had no significant differences in blood or adipose tissue parasitemia compared to infected wild type mice (Figure 6C-D). Finally, to determine the importance of parasite resistance and adipose tissue wasting for outcome of infection, we measured survival in our different genetic models. Both Rag1 KO and B cell KO mice were highly susceptible to *T. brucei* infection, with 100% of mice succumbing to the infection by ~20 days both infection, demonstrating the critical role for resistance defenses in survival of the infection (Figure 6G). CD8 KO mice were protected from infection induced lethality, exhibiting an ~25% increase in the median time to death compared to wild type mice, suggesting that CD8+ T cells compromise the ability of the host to tolerate the infection, likely by contributing to immunopathology (Figure 6G). CD4 KO and *Pnpla2 Cre+* mice exhibited similar death kinetics as their respective wild type controls demonstrating that sickness induced anorexia and adipose tissue wasting have no influence on host outcome of *T. brucei* infection (Figure 6G-H). Taken together, our data demonstrate a decoupling of adaptive immune mediated resistance mechanisms that control parasitemia from the adaptive immune system-mediated metabolic responses to infection and that the adipose tissue wasting has no impact on tissue inflammation, the humoral response or outcome of a chronic parasitic infection (Figure 6I).

**Figure 6.**
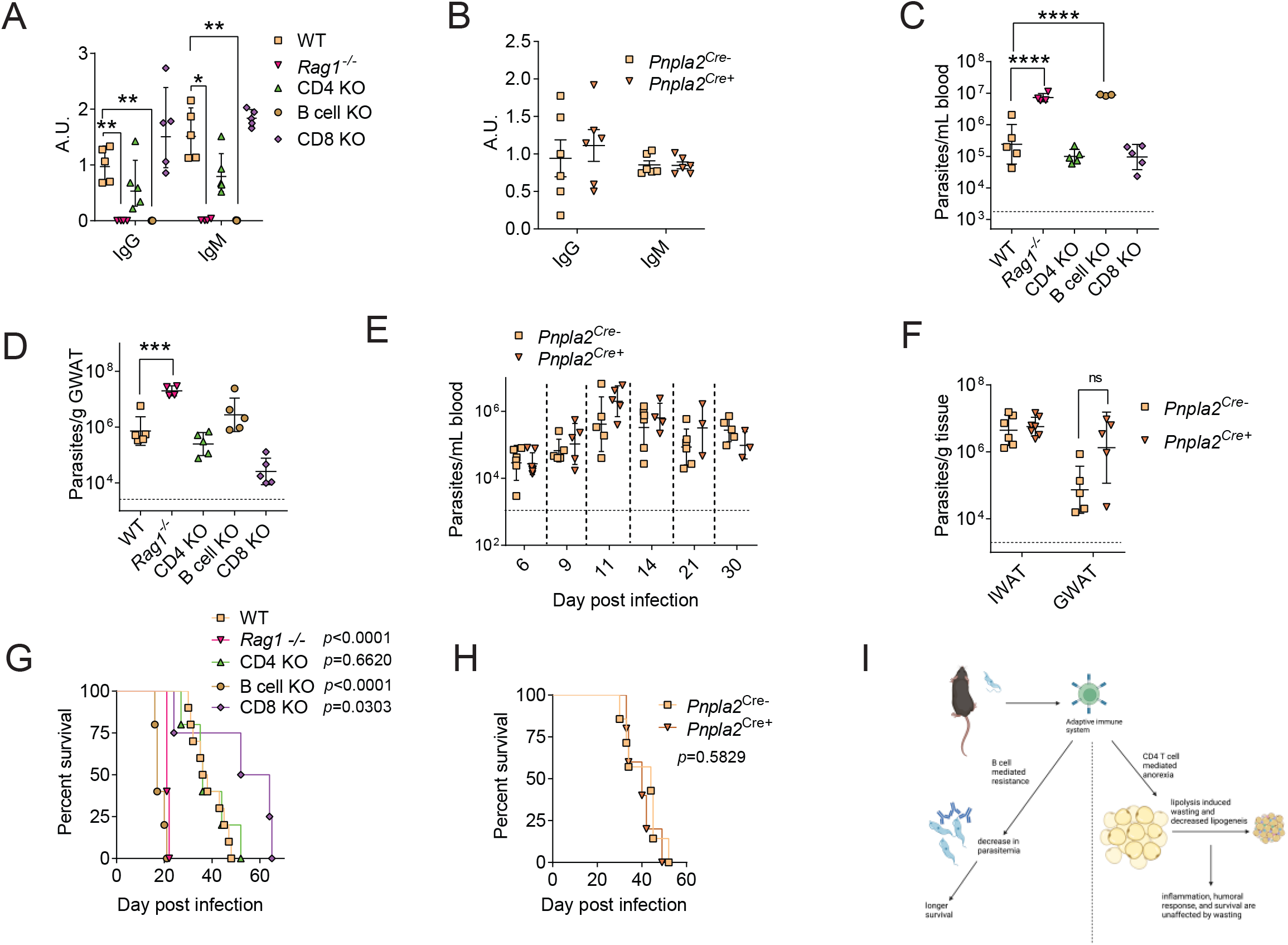
Decoupling resistance from adipose tissue wasting during *T. brucei* infection. (A) WT and immune cell deficient mice were infected with 500,000 *T. brucei*. Antibodies in the serum of infected WT (n=5), *Rag1* -/- mice (n=5), CD4 KO (n=5), CD8 KO (n=5), and B cell KO (n=5) that are specific to *T. brucei* determined via ELISA displayed in arbitrary units. IgG was a 1/25 dilution of serum while the IgM was a 1/200 dilution of serum. These data are taken from the titrations shown in Supplemental Figure 9A-B. (B) Antibodies in the serum of infected *Pnpla2^Cre+^* (n=6) and *Pnpla2^Cre-^* littermates (n=6) that are specific to *T. brucei* displayed in arbitrary units at 7 days post-infection. Serum was diluted 1/1000 for both antibody isotypes. (C) Blood parasitemia at 7 days post-infection of infected WT (n=10), *Rag1* -/- (n=5), CD4 KO (n=5), CD8 KO (n=5), and B cell KO (n=5). The limit of detection was 1350 parasites/mL, represented by a dotted line on the graph. (D) Parasitemia of gonadal white adipose tissue (GWAT) of infected WT (n=5), *Rag1* -/- (n=4), CD4 KO (n=5), CD8 KO (n=5), and B cell KO (n=5). The limit of detection was 3400 parasites/g of fat, represented by a dotted line on the graph. (E) Parasitemia of blood of infected *Pnpla2^Cre+^* (n=5) and infected *Pnpla2^Cre-^* littermates (n=6). The limit of detection was 1350 parasites/mL, represented by a dotted line on the graph. (F) Parasitemia of both GWAT (n=5 for each group) and inguinal white adipose tissue (IWAT) of infected *Pnpla2^Cre+^* (n=7) and infected *Pnpla2^Cre-^* littermates (n=6). The limit of detection is 3400 parasites/g of fat and is represented by a dotted line on the graph. (G) Survival curve of infected C57BL/6J (WT) (n=10), B6.129S7-Rag1^tm1Mom^/J mice (*Rag1* -/-) (n=5), B6.129S2-Cd4t^m1Mak^/J (CD4 KO) (n=5), B6.129S2-Cd8a^tm1Mak^/J (CD8 KO) (n=5), and B6.129S2-Ighm^tm1Cgn^/J (B cell KO) (n=5). (H) Survival curve of infected *Pnpla2^flox/flox^;Fabp4^Cre+^* (*Pnpla2^Cre+^*) (n=11) mice and *Pnpla2^flox/flox^;Fabp4^Cre-^* (*Pnpla2^Cre-^*) littermates (n=7). All infections were carried out with an intravenous injection of 500,000 parasites. * p<0.05, ** p<0.01, *** p<0.001, and *** p<0.0001 Survival curves were analyzed with Logrank (Mantel-Cox) test. For pairwise comparisons, One-way ANOVA comparing all means to infected B6 mice using the Tukey test for multiple comparisons or Kruskal Wallis for non parametric tests. Error bars represent +/- SEM except in panels C, D, E, and F which are geometric mean plotted with geometric SD as the error bars. One representative experiment shown for each panel.

## Discussion

Catabolic processes that occur during an infection are intimately linked to the host immune response to the pathogen. While a role for inflammatory cytokines in driving these catabolic processes is well appreciated, the immune cells that mediate this metabolic response and whether it serves any role for host defense remains poorly understood. Here, using a mouse model of a chronic parasitic infection, we demonstrate that CD4+ T cells are a driver of the infection induced anorexic response, which results in the induction of lipolysis in adipose tissue and fat wasting. We demonstrate that this wasting is dependent on ATGL activity in adipose tissue that is activated in response to reduced food consumption. Surprisingly, we found that while the adaptive immune response drives adipose tissue wasting, this metabolic response serves no role in host resistance defenses against the parasite or the outcome of infection. While catabolic processes have been postulated to be necessary to fuel the host resistance response to infection, our data demonstrate a decoupling of resistance defenses and lipid mobilization. Our findings highlight the diversity of mechanisms through which the host immune system can trigger a common metabolic response during an infection.

In agreement with previous reports (Trindade et al., 2016), we found that *T. brucei* localizes to the white adipose tissue of the infected host. One possible explanation for why adipose tissue can harbor parasites is that the tissue does not have a strong immune response to infection allowing the tissue to become a reservoir of the pathogen (Tanowitz et al., 2017). From our immune phenotyping of white adipose tissue, we found that, consistent with previous reports (Machado et al., 2021) that the adaptive immune response is activated in the white adipose tissue early in the infection course and that this correlates with the decline of parasite burdens in the fat. While we found *T. brucei* infection to increase the levels of activated CD4+ and CD8+ T cells in the adipose tissue, with no change in B cell numbers or activation in the fat during infection, our results with cell specific knock out animals clearly demonstrates that B cells and not CD4+ or CD8+ T cells are necessary for promoting resistance defenses in the white adipose tissue. This indicates that the localization is not due to the fat being immune privileged and that the immune response is capable of decreasing the parasitemia within the tissue, which corresponds to other research indicating that the localization of the parasite may be due to other factors (Machado et al., 2021; De Niz et al., 2021). Further research into why *T. brucei* and other parasites localize to fat and whether it serves any benefit to the pathogen is needed.

In models of cancer cachexia, lipolysis is necessary for the depletion of fat stores, however in a study examining adipose tissue wasting during a chronic viral infection it was suggested that the wasting occurs independent of lipolysis (Baazim et al., 2019) (Daas et al., 2018b; Dahlman et al., 2010; Das et al., 2011; Fukawa et al., 2016; Zagani et al., 2015). In our study, we found that adipose tissue wasting is dependent on *Pnpla2* specifically in adipocytes and occurs downstream of the anorexic response. The simplest explanation to explain why Baazim et al did not observe lipolysis dependent wasting with lymphocytic choriomeningitis virus (LCMV) but we did in our study is that this can reflect differences in viral versus parasitic infections. A second explanation could be due to technical reasons. Baazim et al. used total body weight loss as their readout for adipose tissue wasting rather than actually measuring body fat of mice deficient for ATGL or HSL specifically in adipocytes infected with LCMV. It was previously reported that mice deficient for ATGL in adipocytes undergo compensatory muscle loss when subjected to conditions that cause negative energy balance (Romero et al. 2022). Therefore, it is possible that lipolysis is required for LCMV induced fat wasting and that the lack of changes in total body weight loss observed by Baazim et al. was due to a compensatory loss of lean energy stores.

CD4+ T cells have been shown in previous studies of parasite and cancer cachexia to be protective against the pathogenesis of cachexia. In the present study, we demonstrate that CD4+ T cells are a driver of adipose tissue wasting in response to a *T. brucei* infection. We further demonstrate that CD4+ T cells are required for the induction of the sickness-induced anorexic response. While our efforts to override the anorexic response with force feeding were not effective because it resulted in the mice eating even less food during the infection, using our pairwise feeding strategy we were able to demonstrate that reduced food consumption was sufficient to induced adipose tissue wasting in *T. brucei* infected mice deficient for CD4+ T cells. This is in contrast to previous studies of viral and cancer induced cachexia that suggested the cachectic response and adipose tissue wasting could not be recapitulated by calorie depletion (Baazim et al., 2019; Ebadi and Mazurak, 2014). We currently do not know what subset of CD4+ T cells are responsible for *T. brucei* induced fat wasting or the molecular mechanism by which these cells regulate the sickness induced anorexic response during the infection. However, we found that CD4+ T cells in the adipose tissue were predominately IFNγ positive with only 1-2% of the CD4^+^ T cells producing IL-4 or IL-17 indicating that there is a strong Th1 response and not a strong Th17 or Th2 response. In an LCMV mouse model, it was shown that CD4+ T cells were dispensable and CD8+ T cells were necessary for adipose tissue wasting (Baazim et al., 2019). Together, these studies demonstrate that a common metabolic response to a diverse array of pathogens can by highly specific. Interestingly a previous study using a parasitic nematode demonstrated that anorexic was also dependent on CD4+ T cells but that this may be dependent on a Th2 response suggesting there may be multiple pathways for CD4^+^ T cells to induce anorexia (Worthington et al., 2013).

An outstanding question in infection biology is why does wasting of energy stores occur. Such catabolic process could benefit the pathogen, optimize host defense, or serve no beneficial function for the host or the pathogen. Adipose tissue wasting could impact infection disease outcome in numerous ways. Fat metabolism is thought to play a role in cell function with specific lipid species either increasing or decreasing the inflammatory state of both myeloid and T cells (Hubler and Kennedy, 2016). In addition, mounting an immune response is thought to be energetically costly and adipose tissue wasting could be a mechanism for lipid mobilization to provide energy for the immune response (Eraud et al., 2005; Ganeshan et al., 2019; Johnson et al., 2021). We found that adipose tissue wasting had no influence on the induction of the immune response in the adipose tissue or spleen in response to a *T. brucei* infection. We further found that adipose tissue wasting had no influence on host resistance to the parasite, antibody response or the outcome of infection. Thus, at least in a murine model of *T. brucei* infection, adipose tissue wasting serves no apparent beneficial function for the host or the pathogen.

Previous work from our lab has demonstrated that during a gut infection with *Salmonella* Typhimurium, anorexia limits pathogen transmission (Rao et al., 2017). *T. brucei* is spread through bites from the tsetse fly and the skin has been reported as an important reservoir of the parasite for transmission (Capewell et al., 2016). We did not evaluate if anorexia, and the adipose tissue wasting it causes, has any impact on subcutaneous fat or parasitemia in the skin. It is tempting to speculate that while we show that anorexia and wasting serves no beneficial role at the individual host level, it could theoretically provide benefits at the group or population level to reduce pathogen transmission. For example, the anorexic response may alter the abundance or physiology of the subcutaneous fat, which may influence the feeding of the tsetse fly and ingestion of parasites. In summary, this work defines a new mechanism by which the adaptive immune system regulates sickness induced anorexia and the resulting adipose tissue wasting and decouples immune driven metabolic responses from host resistance mechanisms.

## Acknowledgments

We are grateful to Cody Fine, Mateo Espinoza, and Ashley Rider (UCSD HESCCF) for technical assistance with flow cytometry experiments. This work was additionally made possible by the UC San Diego Stem Cell Program and a CIRM Major Facilities grant (FA1-00607) to the Sanford Consortium for Regenerative Medicine. This publication includes data generated at the UCSD Human Embryonic Stem Cell Core Facility. Flow Cytometry data was generated using the BD Biosciences Fortessa X20 HTS analyzer. This work was supported by NIH awards DPI AI144249 and R01AI14929, and the NOMIS Foundation (J.S.A.). This work was supported by the Flow Cytometry Core Facility of the Salk Institute with funding from NIH-NCI CCSG: P30 CA014195. Figure 6 utilized images from BioRender. We thank Susan Kaech (Salk Institute) for providing flow cytometry reagents.

## Author Contributions

SER- Conceptualization, Experimental design, data collection, data analysis, writing manuscript SKV- Experimental design, Data analysis

KKS – Data collection

JSA- Conceptualization, experimental design, data analysis, writing manuscript

## Declaration of Interests

JSA holds an adjunct position of UC San Diego and is a member of the Rainin Foundation Advisory Board. The authors declare no conflicts of interest.

## Material and Methods

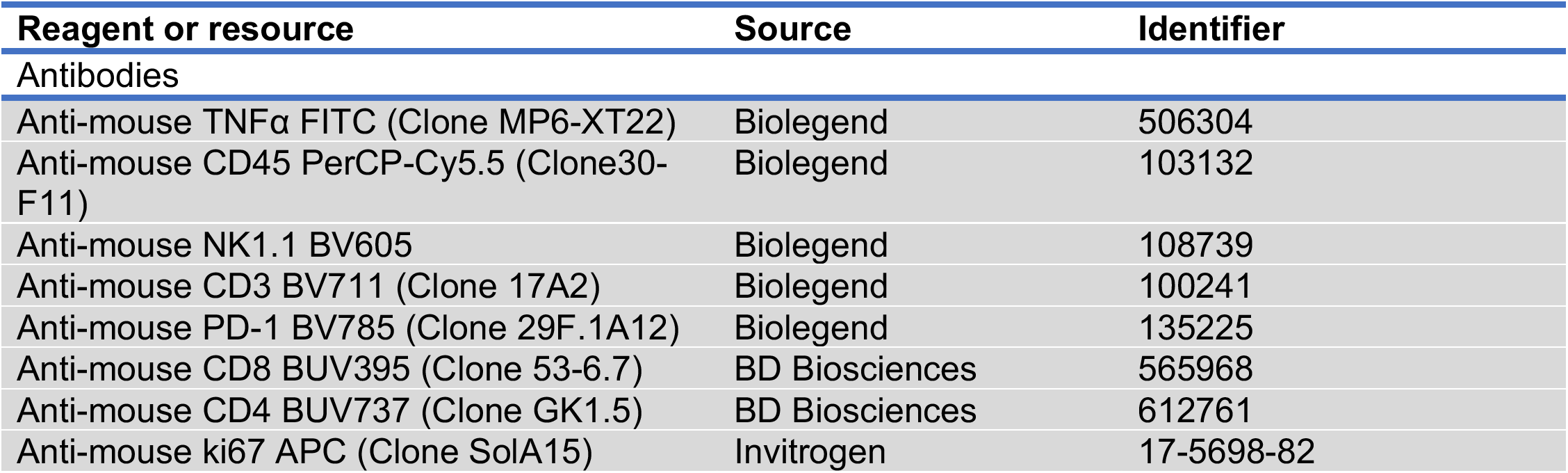
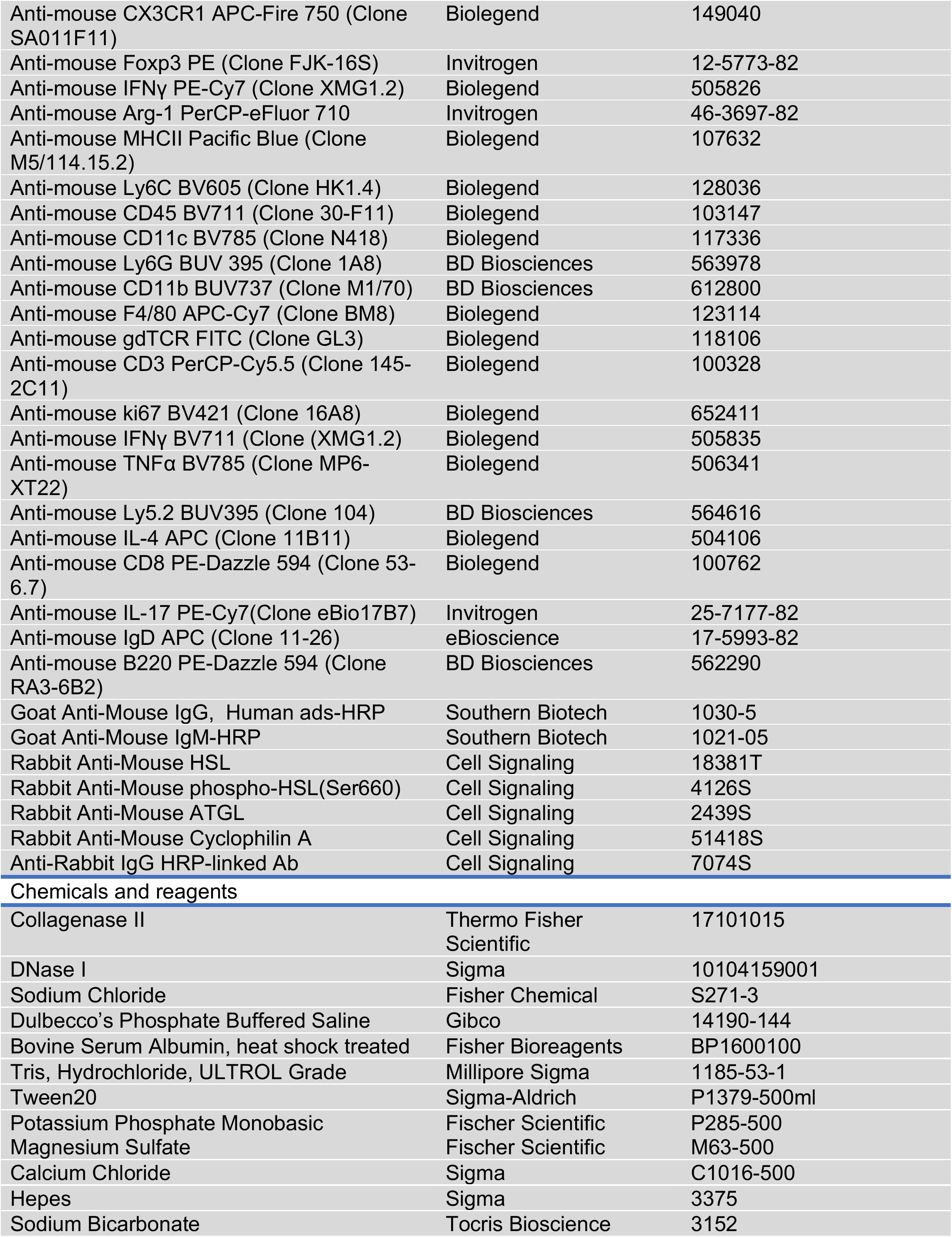
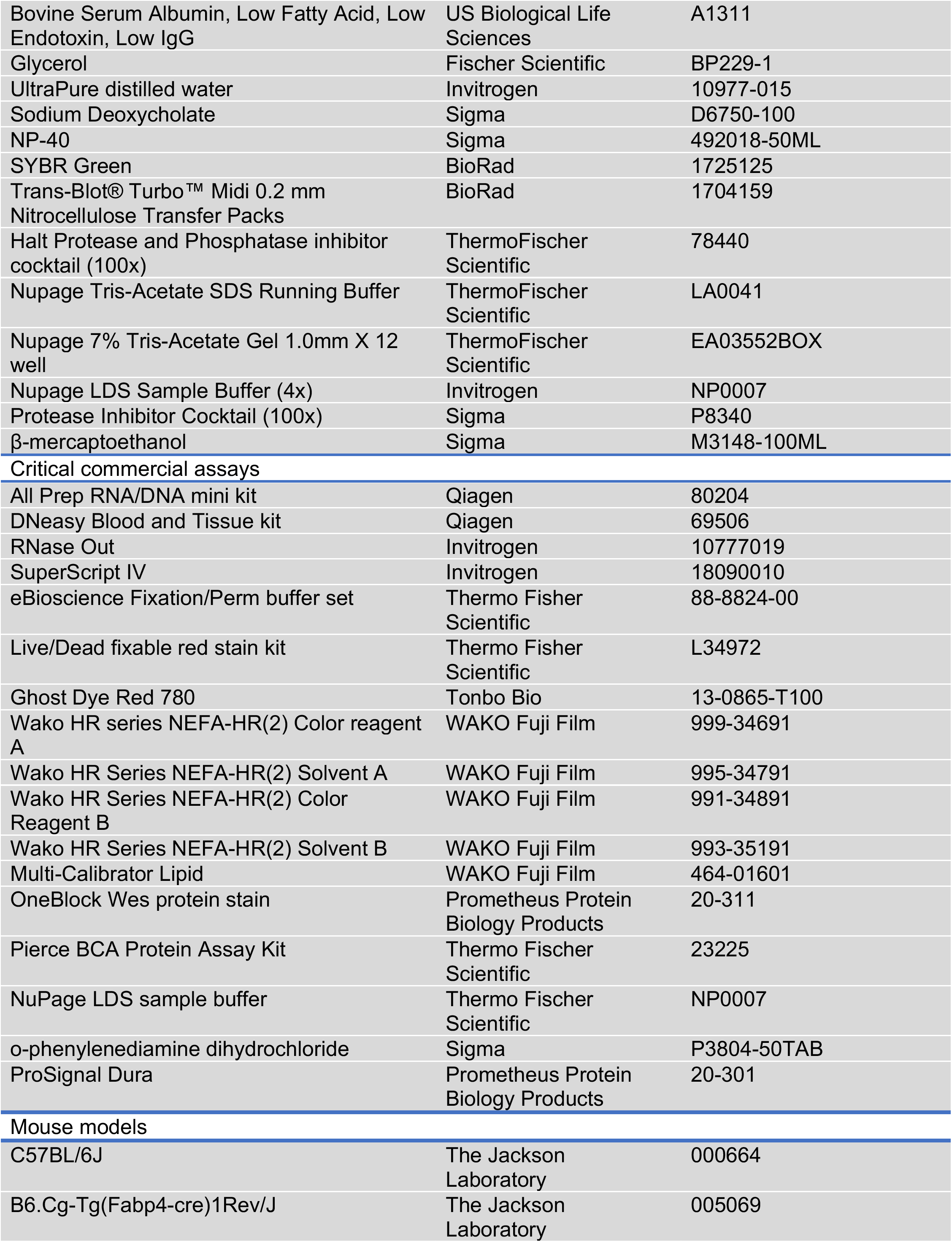
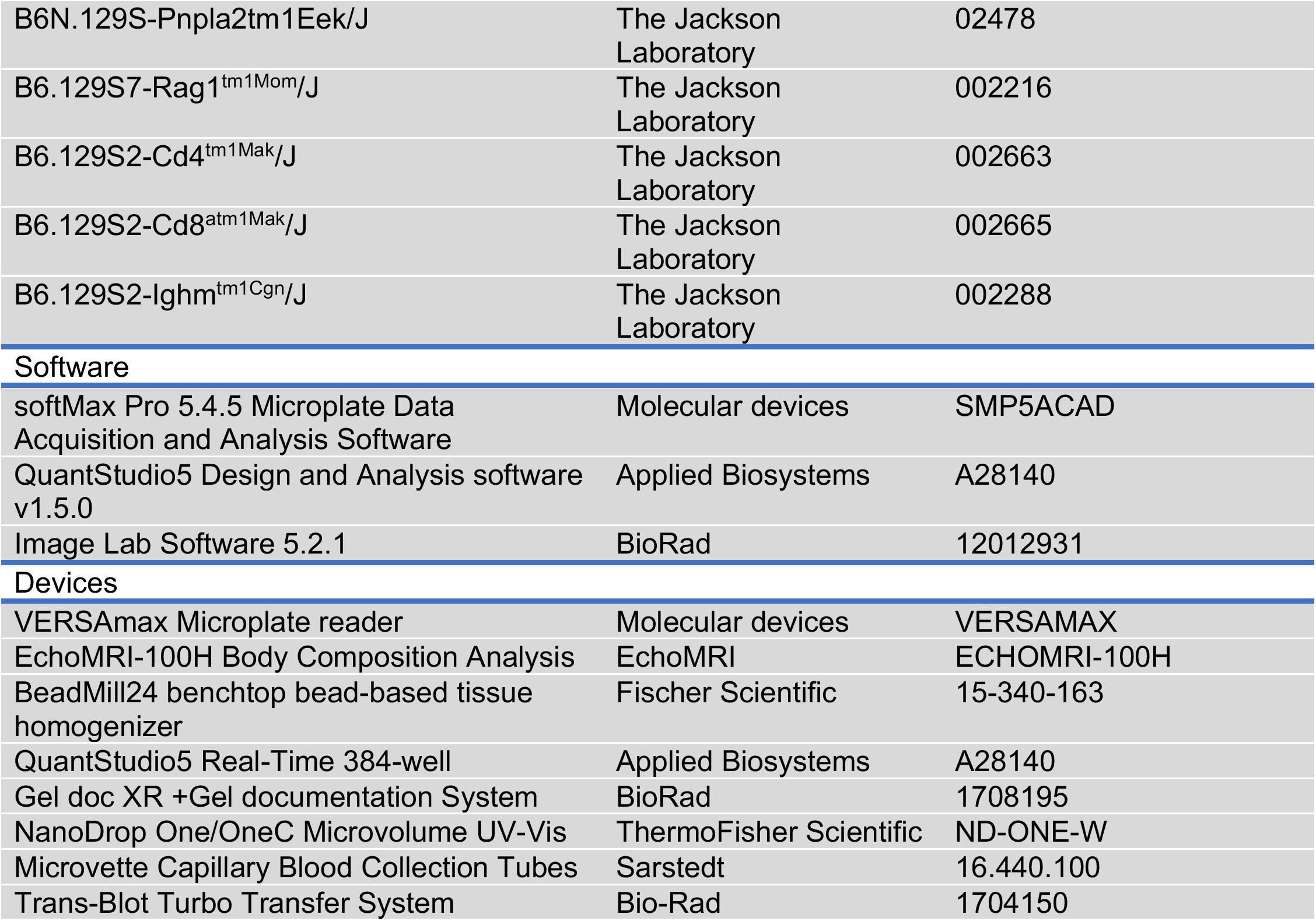

### Mice

Male and female mice of 8-10 weeks of age that were purchased from Jackson Labs or bred in our AALAC-certified vivarium were used for the studies described. For experiments with C57BL/6 mice, animals were purchased from Jackson Labs and acclimated in our facility prior to experimentation. For experiments with B6.129S7-Rag1^tm1Mom^/J mice, animals were either purchased from Jackson Labs or bred in house and then used for experimentation. For experiments with adaptive immune cell specific knock out mice, B6.129S2-Cd4^tm1Mak^/J mice, B6.129S2-Cd8^atm1Mak^/J mice, and B6.129S2-Ighm^tm1Cgn^/J mice were purchased from Jackson Labs along with wild type controls, housed in our vivarium for a few days to acclimate prior to experimentation. To study ATGL dependent lipolysis, we performed crosses with *B6N.129S-Pnpla2tm1Eek/J* mice to *B6.Cg-Tg(Fabp4-cre)1Rev/J* mice (both from Jackson Labs) to generate *Pnpla2;Fabp4^cre+^* and *Pnpla2;Fabp4^cre-^* mice. For experiments, both male and female mice were used at 8-10 weeks of age. Mice were specific pathogen-free, maintained under a 12-hour light/dark cycle, and given standard chow diet *ad libitum* except in pairwise feeding experiments. All animal experiments were done in accordance with The Salk Institute Animal Care and Use Committee.

### Parasites

*Trypanosoma brucei* subspecies *brucei* (Antat1.1) was a gift from Samuel J. Black (University of Massachusetts Amherst).

### Mouse infection models

Mice were intravenously infected with 500,000 moving *Trypanosoma brucei brucei* strain Antat1.1 parasites suspended in PBS that were diluted from frozen stocks. For the preparation of parasite stocks, C57BL/6J mice were irradiated with 950 rads for ~12 minutes. The mice were then interperitoneally injected with parasite stocks provided by Dr. Samuel Black (University of Massachusetts Amherst). A blood smear was obtained daily by tail bleed and parasitemia was quantified using a hemocytometer. Once levels reached ~5*10^7^ parasites per ml of blood, mice were then euthanized, and blood was collected via cardiac puncture with a heparinized needle. The blood was mixed 1:1 with ice cold PBS+20% glycerol. The mixture was then aliquoted into cryotubes with 125 μL of the mixture. Aliquots were placed in a −80°C freezer overnight before being moved to liquid nitrogen storage.

### Culturing *T. brucei*

*Trypanosoma brucei* was initially cultured by adding 50 μL of infected mouse blood to 1 mL of HMI-9 medium (recipe below). The culture was incubated at 37°C and 5% CO2 in a tissue culture incubator. The culture was maintained at 10^5^ - 10^6^ parasites/mL for the first week of culturing. Parasites were then grown to 10^6^ to 10^7^ parasites/mL of culture. They were then centrifuged at 800g for 5 min and resuspended in fresh media at 10^7^ parasites/mL. Parasites were then mixed in a 1:1 ratio with new media plus 20% glycerol and aliquoted into cryotubes and frozen. These cryotubes of culture-adopted parasites were used for growing parasites to coat ELISA plates for antibody quantification.

#### HMI-9 media

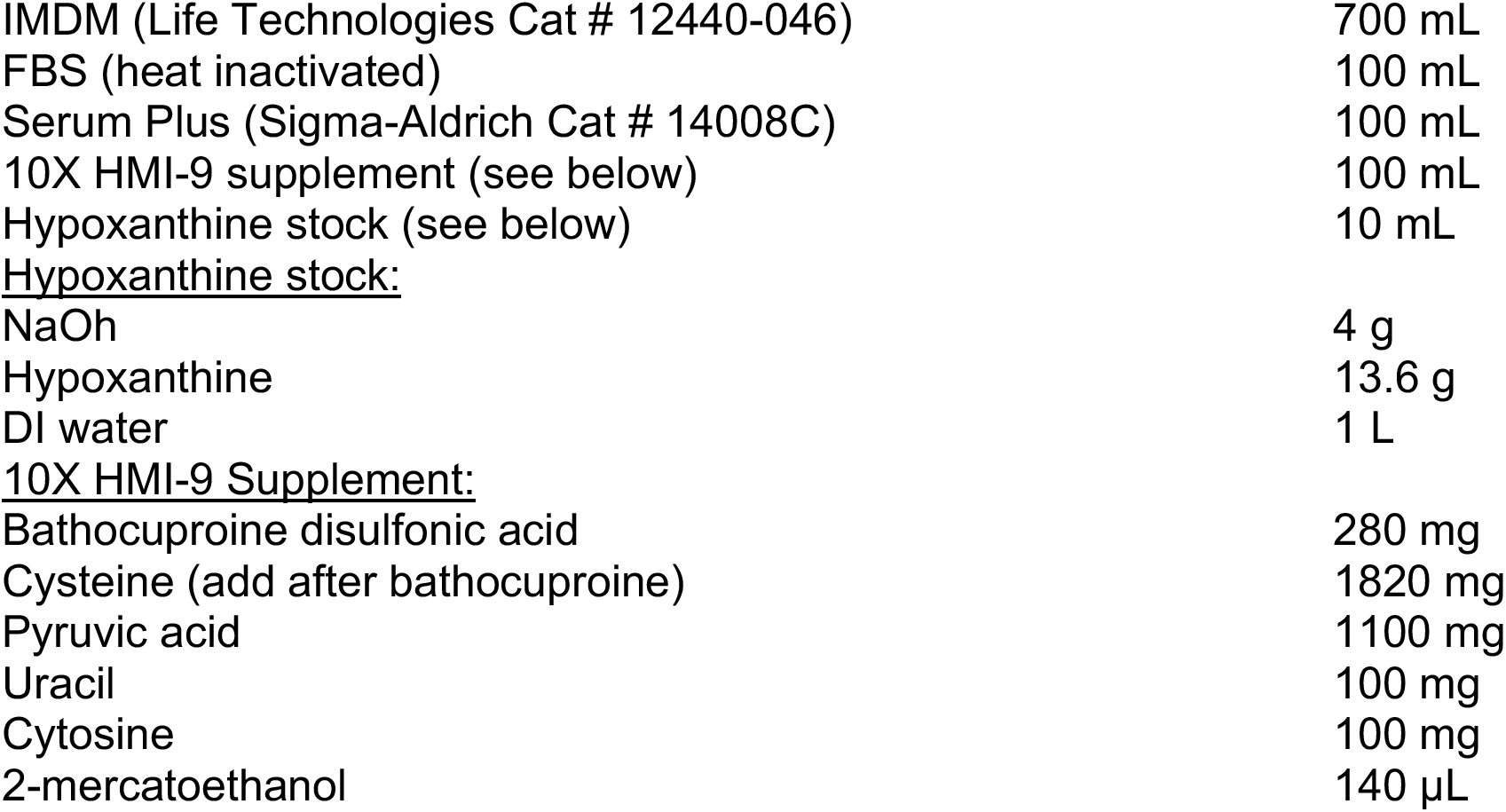

**HMI-9 media:**
|  |  |
| --- | --- |
| IMDM (Life Technologies Cat # 12440-046) | 700 mL |
| FBS (heat inactivated) | 100 mL |
| Serum Plus (Sigma-Aldrich Cat # 14008C) | 100 mL |
| 10X HMI-9 supplement (see below) | 100 mL |
| Hypoxanthine stock (see below) | 10 mL |
| **Hypoxanthine stock:** |  |
| NaOH | 4 g |
| Hypoxanthine | 13.6 g |
| DI water | 1 L |
| **10X HMI-9 Supplement:** |  |
| Bathocuproine disulfonic acid | 280 mg |
| Cysteine (add after bathocuproine) | 1820 mg |
| Pyruvic acid | 1100 mg |
| Uracil | 100 mg |
| Cytosine | 100 mg |
| 2-mercaptoethanol | 140 μL |

### DNA isolation for parasite quantification

DNA was isolated from blood and tissue for parasite quantification using the Qiagen DNeasy Blood and Tissue kit following manufacturer’s protocol. Briefly, either 25 μL of heparinized blood or 10-20 mg of tissue were placed in 180 μL ATL buffer with 20 μl of proteinase K at 56°C until blood or tissue was lysed. 200 μL of AL buffer was added and the samples were mixed. 200 μL of 100% ethanol was then added and the samples were mixed again. The samples were then loaded onto a spin column and centrifuged at 6000 rpm for 1 minute. The supernatant was discarded, and the column was washed with 500 μL of AW1 buffer before the sample was centrifuged again at 6000 rpm for 1 minute. The column was then washed with 500 μL of AW2 and the column was centrifuged for 3 minutes at max speed to dry the column. The column was placed into a new tube and 200 μL of AE buffer was added and the column was incubated for a minute before being spun at 6000 rpm for 1 minute to elute the DNA.

### Parasitemia via qPCR

To determine parasitemia in the blood and tissue, a standard curve was generated by culturing *T. brucei to* 1.36*10^7^ parasites per mL. DNA was extracted from 25 μL of culture using Qiagen DNeasy Blood and Tissue kit following the manufacturer’s protocol. Serial dilutions were made from the DNA \qPCR was done using forward primer (ACGGAATGGCACCACAAGAC) and reverse primer (GTCCGTTGACGGAATCAACC) (Trindade et al., 2016) to generate a standard curve. DNA from the blood or tissue of mice was extracted using the same kit (Qiagen blood and tissue kit) and qPCR was done using the same primers. The Ct of the extracted samples were then compared to the standard curve to determine the number of parasites in the sample.

### Body composition measurements

Total body fat and lean mass were measured using an EchoMRI machine. The total fat and total lean mass readings were normalized to the weight of the individual mouse at the start of the experiment. Fat pad masses were measured by dissecting and weighing the subcutaneous fat pad (inguinal white adipose tissue [IWAT]) and the visceral white adipose tissue (gonadal white adipose tissue [GWAT]). The fat pad weights were normalized to the starting weight of the mouse. The gastrocnemius was dissected and weighed as a representative of muscle tissue and normalized to starting weight of the mouse.

### Protein isolation

Frozen tissue was ground into a powder and treated with 900 μL of lysis buffer (50 mM Tris pH 7.5, 150 mM NaCl, 1% NP-40, 0.5% Sodium Deoxycholate, 5% Glycerol, 0.1% sds, 1mM EDTA [pH 8]) along with Halt phosphatase inhibitor and sigma protease inhibitor cocktail. Tissues were homogenized by using a ceramic bead and a BeadMill24 shaken at 3200 rpm for 30s and were immediately placed on ice. The samples were centrifuged at max speed for 10 minutes and the middle layer was collected. The sample was then spun again at max speed for 30 minutes to take the middle layer again. Finally, the samples were spun at max speed for 40 minutes and the top layer was removed by using a pipette to try to remove any residual lipids. The sample was then placed on ice and protein concentration was measured using BCA kit.

### Western blot

For western blots, 60 μg of protein suspended in 60 μL of lysis buffer was mixed with 15 μL of LDS/BME. Samples were then incubated at 95°C for 10 minutes before sonicating for 10 minutes. 15 μL of sample was then loaded onto a NuPage 7% Tris acetate gel. The same lysates were used to run the blots for total and phospho HSL and the corresponding loading controls. Gels were run for 150V for 60 minutes in an Invitrogen Mini Gel Tank and transferred to nitrocellulose blot using the Bio-Rad Turbo Transblot system for 10 minutes at 25V (1.3A). Staining was done with OneBlock Wess protein stain which was used to cut the membrane into strips according to the size of the protein target. These strips were washed in TBST (20mM Tris, 150mM NaCl, 0.1% Tween20 w/v) and blocked with 5% BSA in TBST for one hour at room temperature. Blots were incubated with the primary antibody overnight (see table above for catalog numbers) at 4°C. Blots were washed the next day, 4 times for 5 minutes each with TBST. They were then incubated for 1 hour at room temperature with Anti-Rabbit IgG HRP-linked antibodies with gentle shaking. They were then washed 4 times for 5 minutes each with TBST. Blots were developed using ProSignal Dura chemiluminescence and imaged in Bio-Rad Gel Doc XR system. Phosphorylated target proteins were probed simultaneously as total protein by using one gel for total and one for phosphorylated targets loaded with the same amount of protein.

### Lipolysis assay

~0.025g of adipose tissue was removed from both the inguinal and gonadal white adipose tissue in duplicate. The tissue was placed in ice cold PBS until samples from all mice were gathered. They were then transferred to 300 μL of Krebss-Ringer bicarbonate hepes buffer (120 mM NaCl, 4 mM KH2PO4, 1mM MgSO4, 0.75 mM CaCl2, 30mM Hepes, 10 mM NaHCO3, 2% fatty acid free BSA, 5mM glucose) and incubated in tissue culture incubators at 37°C and 5% CO2 for 4 hours. The supernatant was then collected and frozen for downstream quantification of free fatty acids and glycerol.

### Free fatty acid measurements

Free fatty acids were measured using the FujiFilm Wako Chemical’s NEFA assay kit following the manufacturer’s protocol. Briefly, 10 μL of supernatant from the lipolysis assay were placed onto a 96 well plate in duplicate. Standards were made by using the 1 mM standard available from FujiFilm Wako and doing ½ serial dilutions to 0.0625 mM concentration of free fatty acids and adding 10 μL of each dilution to the 96 well plate in duplicate. Then, 135 μL of reagent A was added to each well with sample or standard in it and the plate was incubated at 37°C for 15 minutes. After incubation, 65 μL of reagent B was added to each well with sample or standard and the plate was incubated at 37°C for 10 minutes. The plate was then read at 550 nM and analyzed according to the manufacturer’s protocols. Readings were taken using a 96-well VERSAmax microplate reader and SoftMax Pro software.

### Glycerol measurements

Glycerol was measured by using Sigma’s Free Glycerol Reagent (F6428). 4 μL of supernatant from lipolysis assay was loaded onto a 96 well plate in duplicate. Standards were made by diluting glycerol to 8, 7,6, 5,4, 3, 2, and 1 mM concentrations and loading 4 μL of each onto plate in duplicate. 150 μL of the Free Glycerol Reagent was added to each well and the plate was incubated at room temperature for 15 minutes. The plate was then read at 540 nM wavelength. Analysis was done by fitting the standards to a linear line and using the equation of that line to calculate the concentration of each sample. Readings were taken using a 96-well VERSAmax microplate reader and SoftMax Pro software.

### Pairwise feeding experiments

Mice were single housed for two days before the beginning of the experiment for acclimation. The infected mice were fed *ad libitum*. The uninfected mice were broken up into two groups, both of which were given water *ad libitum*. One group of uninfected mice was given food *ad libitum* while the other group was given the same amount of food that an average infected mouse ate each day during a previous experiment (shown in Figure 5A). Food was given at 10 am each morning for that day’s food intake. At 0, 7- and 13-days post-infection, mice were dissected to determine their fat pad weights.

### qPCR of lipogenesis genes

RNA was isolated from bead-beaten inguinal white adipose tissue using Qiagen All Prep RNA/DNA Mini Kit. Briefly, the sample was lysed in 600 μL RLT buffer containing β-mercaptoethanol and centrifuged for 3 minutes at 6,000 rpm in a microcentrifuge. The supernatant was transferred to an AllPrep DNA column and centrifuged for 30 s at 10,000 rpm. Flow-through was mixed with 70% ethanol and transferred to an RNeasy spin column and centrifuged for 30 seconds. The flow-through was discarded and the column containing RNA was washed with 350 μL RW1 buffer. Flow-through was discarded. The column was treated with 80 μL of Qiagen RNase-free DNase1 for 15 minutes at room temperature. The column was washed again with 350 μL RW1 and the flow-through was discarded. The column was then washed with 500 μL of RPE buffer. After the final wash, the column was placed in a new Eppendorf tube, and RNA was eluted with 40 μl of RNase-free water. cDNA was made using the SuperScript IV kit (Invitrogen) after all samples were diluted to the concentration of RNA in the lowest sample. Real-time qPCR was performed using iTaq SYBER Green Mix (BioRad) on a ThermoFisher QuantStudio 5 qPCR machine. Relative standard curves were generated by mixing small amounts of cDNA from each sample and then making serial dilutions of the mix. Analysis of gene expression was then done by comparing Ct of the gene of interest to the relative standard curve for that gene, then normalizing this to the expression of the housekeeping gene for each sample. The housekeeping gene used was *Rps17*. See table 1 for primer sequences.

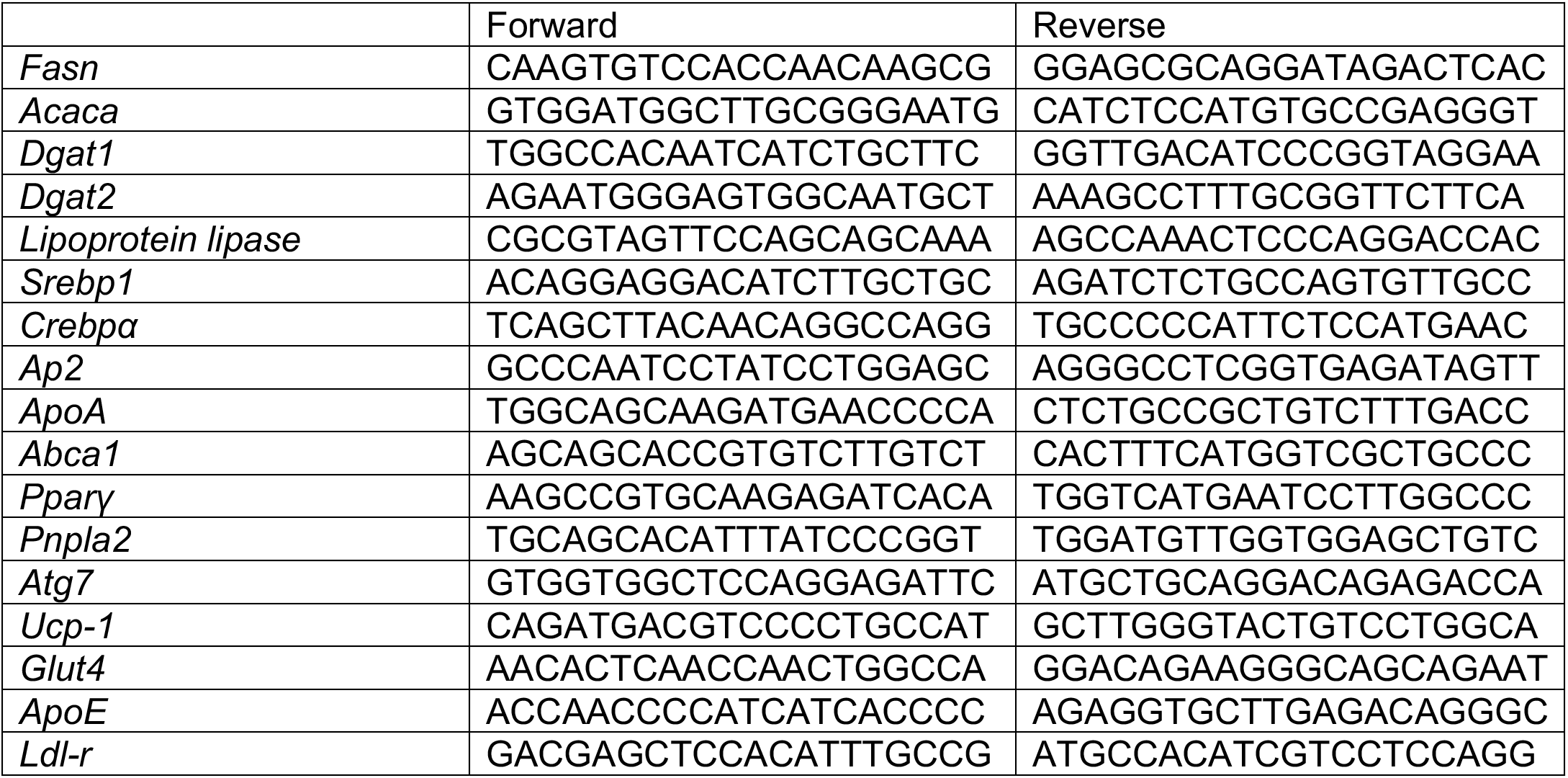

### ELISA for antibody relative quantification

Parasites were grown to 9.8*10^6^ parasites per mL of culture. They were then heat killed at 56°C for one hour before freezing. The day before ELISA was performed, the parasites were thawed and 200 μL of HK parasites were added to each well of a Nunc ELISA plate and incubated at 4°C overnight. The next day, the plate was washed with PBS+0.1% TWEEN. Serum was diluted in PBS+1% Triton+1% BSA and 100 μL of the dilution was added to the plate in duplicate for each isotype being tested. Plate was then incubated at 37°C for one hour before the plate was washed again. A secondary antibody was then added which was either anti-mouse IgG or anti-mouse IgM, both of which were conjugated to HRP. The plate was incubated for another hour at 37°C. The plate was washed again and then developed using o-phenylenediamine dihydrochloride. The reaction was stopped with 20 μL of 3M HCl before being read on a VERSAmax microplate reader using SoftMax Pro software.

### Cell isolation, stimulation, staining

To isolate cells from fat, gonadal white adipose tissue was removed from the mouse, minced with a sterilized razor blade, and placed in ice-cold PBS. The tissue was then treated with 0.5 mg/mL of collagenase II and 0.5 mg/mL of DNAse I for 30 minutes at 37°C shaking at 250 rpm. After digestion, 10 mL of 10% heat inactivated FBS+PBS was added to neutralize the collagenase. The tissue was then passed through a 70 μm filter. The solution was then centrifuged at 400 g for 10 minutes and the supernatant was removed using a vacuum. 500 μL of ACK lysis was added and samples were incubated for 2 minutes before 5 mL of PBS+10% heat inactivated FBS were added to dilute the ACK buffer. The solution was centrifuged at 400g for 10 minutes. Supernatant was then discarded, and the sample was resuspended in 200 uL of FACS buffer and placed onto a 96 well plate. Cells were counted using a hemocytometer. The plate was centrifuged for 3 minutes at 500g. T supernatant was removed, and stimulation reagents were added to the cells. For T cells, golgi plug was added along with PMA and ionomycin at 10 ng/mL. For the myeloid cells, either heat killed *T. brucei* was added at about 10^6^ parasites/mL or LPS was added at 10 ng/mL. The cells were incubated for 4 hours before they were centrifuged at 500 g for 3 minutes and supernatant was removed. Then the cells were treated with FC block and live dead stain for 30 minutes. The cells were then centrifuged at 500g for 3 minutes before incubating with the stains for 30 minutes. The cells were then centrifuged again at 500g for 3 minutes before they were fixed and permeabilized overnight using ThermoFisher Fix & Perm kit following manufacturer’s instructions. The following day the cells were washed with FACS buffer and stained for 30 minutes with intracellular stains. The cells were then washed once again and resuspended in 200 μL of FACS buffer. The cells were then run on a Fortessa X20.

### Quantification and statistical analysis

All statistical tests were carried out using Prism 7.05. Sample sizes, statistical tests used, and p values for each figure and panel are indicated in the figure legends. The data shown are either one representative experiment or individual experiments combined and is noted in the figure legends.

## Figure Legends

**Supplementary Figure 1.**
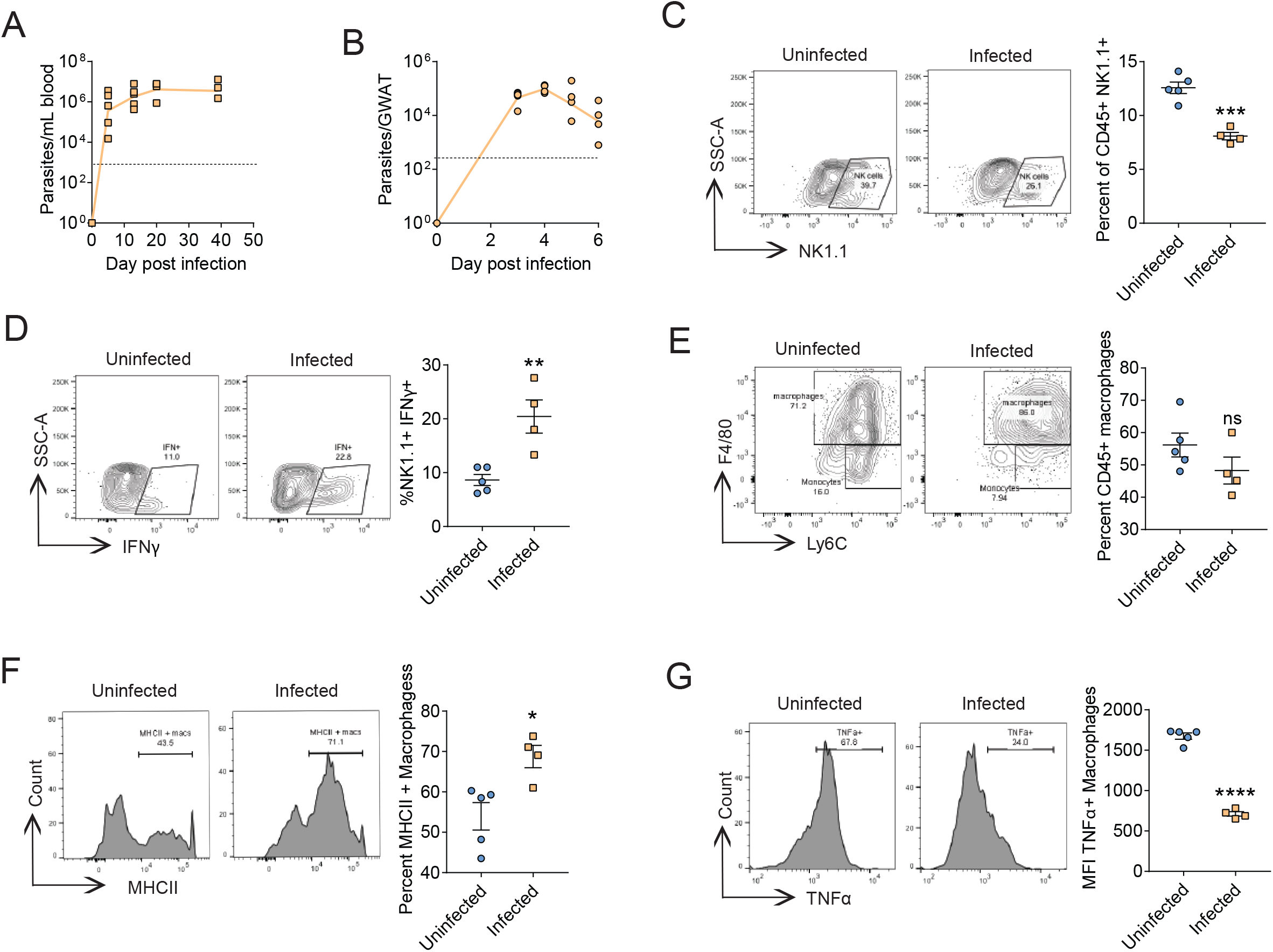
*Trypanosoma brucei* infection activates the adaptive immune response in adipose tissue. 8 week old C57Bl/6 male mice were intravenously infected with 500,000 *T. brucei*. (A) Data from Figure 1B showing individual data points of parasitemia in WT C57BL/6J mice determined via qPCR. n=5 for the first two time-point, n=4 for the last two time-points. The limit of detection was 1350 parasites per mL of blood and is represented by the dotted line on the graph. (B) Data from Figure 1C showing individual data points of parasitemia in the gonadal white adipose tissue (GWAT) of WT C57BL/6J mice determined via qPCR. n=5 for days 3 and 5, n=4 for days 4 and 6. The limit of detection was 3400 parasites per gram of tissue, represented by the dotted line on the graph. (C-G) Additional flow analysis of GWAT from mice displayed in Figure 1 flow panels. (C) Percentage of CD45+ CD3^-^ cells which were NK1.1+ indicating they are natural killer cells. (D) Percentage of natural killer cells from panel C that were IFNγ^+^. (E) Percentage of CD45^+^ CD11b^+^ Ly6G^-^ cells that were macrophages (Ly6Cint-hi, F4/80+). (G) Percent of macrophages from panel E that were MHCII+. (H) Median Fluorescent Intensity of TNFa in macrophages displayed in panel E. * p<0.05, ** p<0.01, *** p<0.001, and **** p<0.0001 unpaired Student’s t-test was done for each panel between infected and uninfected mice. Error bars represent +/- SEM. Frequencies displayed on gates are percent of parent gate which may be different than what is being displayed on graph which is listed above for each panel. Data represent one independent experiment.

**Supplemental Figure 2.**
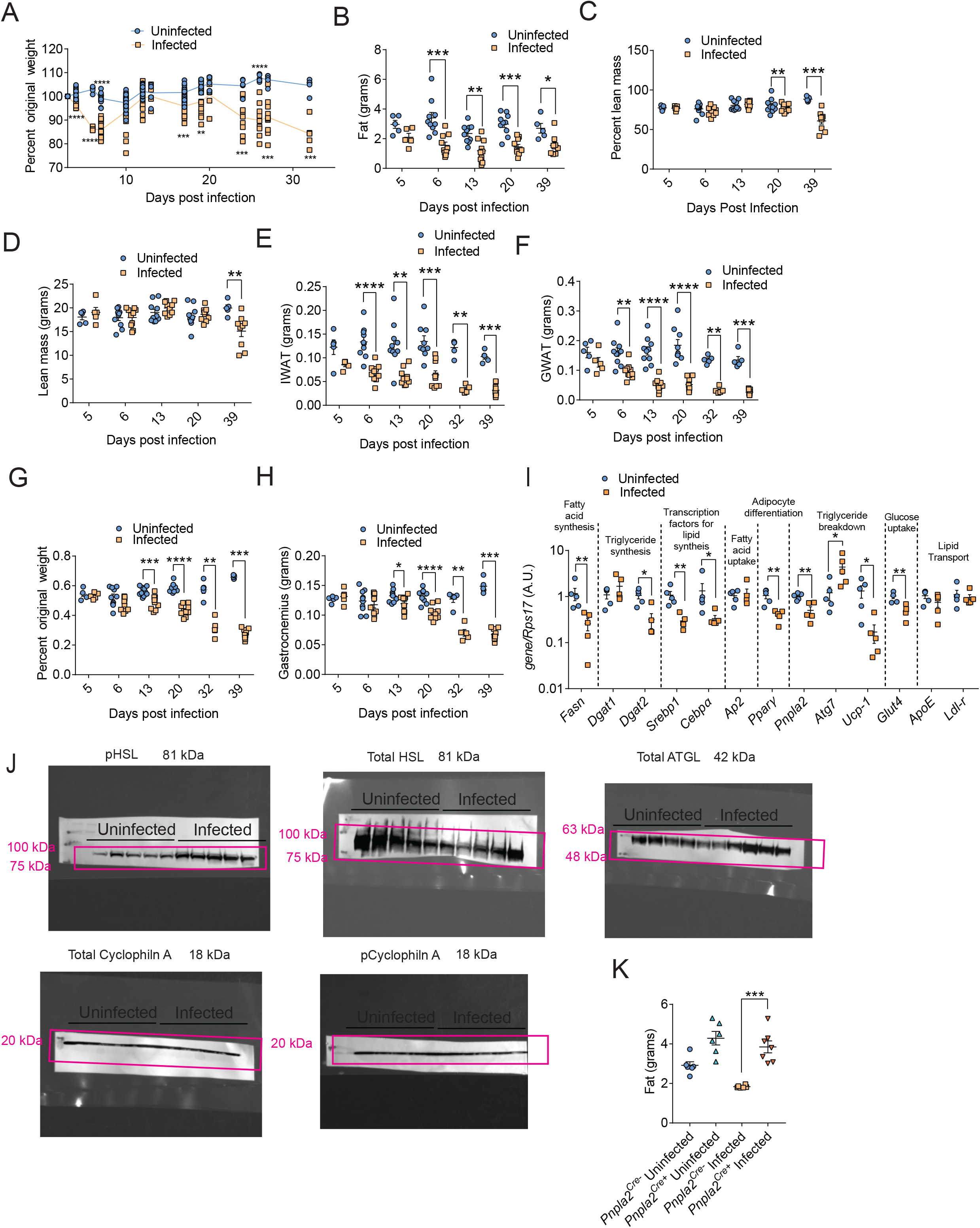
Mice infected with *Trypanosoma brucei* undergo lipolysis-dependent adipose tissue wasting. Mice were intravenously infected with 500,000 *T. brucei*. (A) Data from Figure 2A showing individual data points for mice that were weighed over the course of the infection and normalized to pre-study weight. (B) Data from Figure 2B displaying MRI readings of fat mass in grams for uninfected (n=5 for 5 days post-infection, n=10 for each other timepoint) or infected (n=5 for 5 days postinfection, n=10 for each other timepoint) without normalizing to pre-study weight. (C) Lean mass weight from MRI readings of infected (n=5-10 for each timepoint) or uninfected (n=5-10 for each timepoint) C57BL/6J mice (WT). Same mice as in Figure 2B. Data graphed as a percentage of pre-study weight (D) Lean mass in grams from MRI readings displayed in previous panel without normalizing to pre-study weight. (E) Same data displayed in Figure 2C showing IWAT weights of WT mice (n=5-10 for each time point) or uninfected control mice (n=5-10 for each time point) without normalizing to pre-study weight. (F) Same data displayed in Figure 2D showing GWAT weights of WT mice (n=5-10 for each time point) or uninfected control mice (n=5-10 for each time point) without normalizing to pre-study weight. (G) Gastrocnemius weight of infected WT mice (n=5-10 for each time point) or uninfected control mice (n=5-10 for each time point) plotted as percent pre-study weight of each mouse. (H) Same data displayed in previous panel, showing Gastrocnemius weight of infected WT mice (n=5-10 for each time point) or uninfected control mice (n=5-10 for each time point) without normalizing to pre-study weight. (I) qPCR results of IWAT of lipid trafficking genes in uninfected (n=5) and infected (n=5) C57BL/6J mice normalized to *Rps17* in arbitrary units (A.U.) at 13 days post-infection. (J) Uncropped images of western blot from Figure 2H. Pink boxes highlight where images were cropped to include in Figure 2. Blot labeled “Total Cyclophilin A” is loading control for total HSL and total ATGL. Blot labeled “pCyclophilin A” is loading controls for pHSL blot. Size of each protein is in black on the right while ladder band sizes are given on the left next to the ladder in pink text. (K) MRI of fat tissue mass in female *Pnpla2^flox/flox^;Fabp4^Cre+^* (*Pnpla2^Cre+^*) mice that infected (n=7) or uninfected (n=6) with *Pnpla2^flox/flox^;Fabp4^Cre-^* (*Pnpla2^Cre-^*) littermates either infected (n=6) or uninfected (n=6) displayed in Figure 2I without normalizing to pre-study weight. * p<0.05, ** p<0.01, *** p<0.001, and *** p<0.0001 unpaired t-tests or Mann-Whitney tests were done depending on the normality of the data for each time point or gene between infected and uninfected mice in panels A-I. One-way ANOVAs were performed comparing the mean of each group to every other group with the Tukey’s test correcting for multiple comparisons for panel K. Error bars represent +/- SEM. Panels B-H show two independent experiments combined, all other panels show one representative experiment.

**Supplementary Figure 3.**
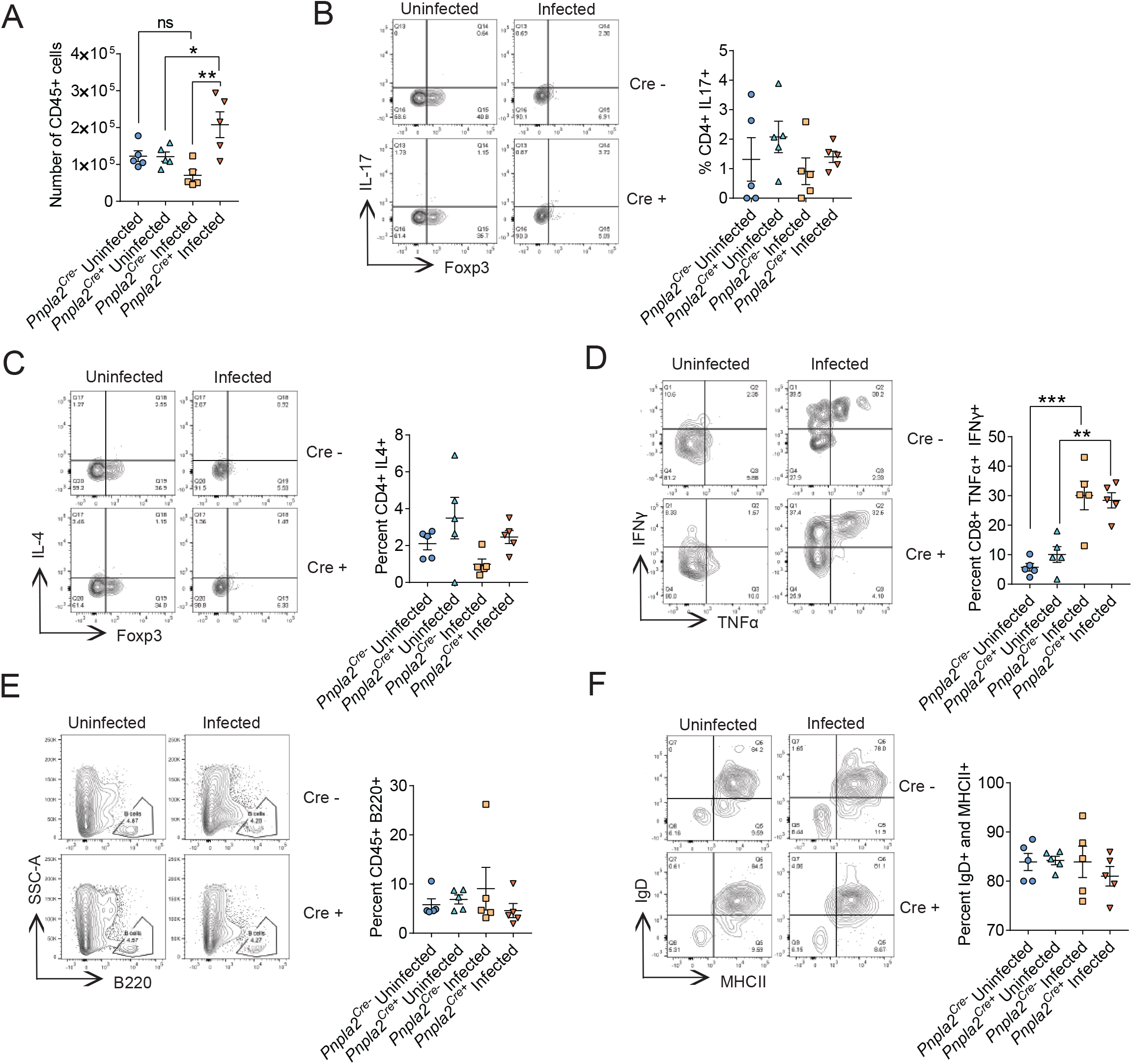
Adipose tissue wasting is not necessary for activation of adaptive immunity in adipose tissue. Mice were intravenously infected with 500,000 *T. brucei*. Cells were isolated from gonadal white adipose tissue (GWAT) of uninfected *Pnpla2^flox/flox^;Fabp4^Cre+^* (*Pnpla2^Cre+^*) mice (n=5), uninfected *Pnpla2^flox/flox^;Fabp4^Cre-^* (*Pnpla2^Cre-^*) (n=5), infected *Pnpla2^Cre-^* (n=5) and infected *Pnpla2^Cre+^* (n=5) littermates at 6 days post-infection and immune cell populations were quantified. These are the same mice as in Fig. 3. (A) Total CD45^+^ cells isolated from the adipose tissue. (B) The proportion of CD4^+^ Foxp3^-^ that are IL-17^+^. (C) The proportion of Foxp3^-^ CD4^+^ T cells that are IL-4^+^. (D) TNFa^+^ IFNγ^+^ CD8^+^ T cells as a proportion of Foxp3^-^ CD8^+^ T cells. (E) The proportion of CD45^+^ cells that were B cells. (F) Percent of B cells that were MHCII^+^ and IgD^+^. * p<0.05, ** p<0.01, *** p<0.001, and *** p<0.0001 One-way ANOVA comparing the mean of each column to every other mean using Tukey correction for multiple comparisons or Kruskal Wallis with a post Dunn test. Error bars represent +/- SEM. Frequencies displayed on gates are percent of parent gate which may be different than what is being displayed on graph which is listed above for each panel. Panels show one representative experiment.

**Supplemental Figure 4.**
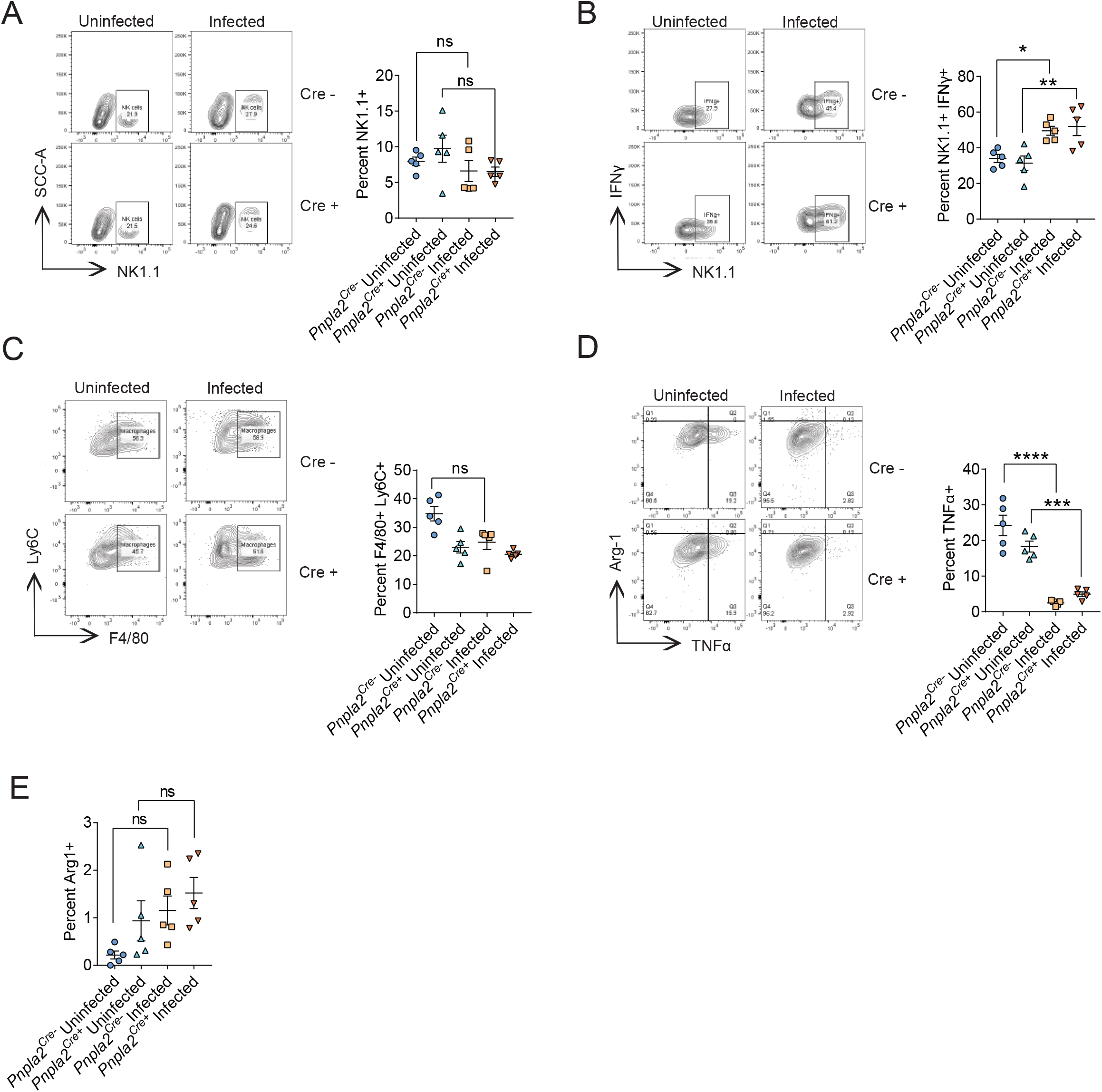
Innate immune cells within the adipose tissue are not affected by adipose tissue wasting. Mice were intravenously infected with 500,000 *T. brucei*. Cells were isolated from gonadal white adipose tissue (GWAT) of uninfected *Pnpla2^flox/flox^;Fabp4^Cre+^* (*Pnpla2^Cre+^*) mice (n=5), uninfected *Pnpla2^flox/flox^;Fabp4^Cre-^* (*Pnpla2^Cre-^*) (n=5), infected *Pnpla2^Cre-^* (n=5) and infected *Pnpla2^Cre+^* (n=5) littermates at 6 days post-infection and immune cell populations were quantified. These are the same mice as in Fig. 3. (A) Percent of CD45^+^ cells that stained with NK1.1 (B) IFNy^+^ Natural Killer cells as a proportion of all Natural Killer cells. (C) Percent of CD45^+^ cells that are macrophages (CD11b^+^, Ly6C^int-hi^, F4/80^+^). (D) Percent of macrophages that are TNFα^+^. (E) Percent of macrophages that are Arg1^+^. * p<0.05, ** p<0.01, *** p<0.001, and *** p<0.0001 One-way ANOVA comparing the mean of each column to every other mean using Tukey correction for multiple comparisons or Kruskal Wallis with a post Dunn test. Error bars represent +/- SEM. Frequencies displayed on gates are percent of parent gate which may be different than what is being displayed on graph which is listed above for each panel. Panels show one representative experiment.

**Supplementary Figure 5.**
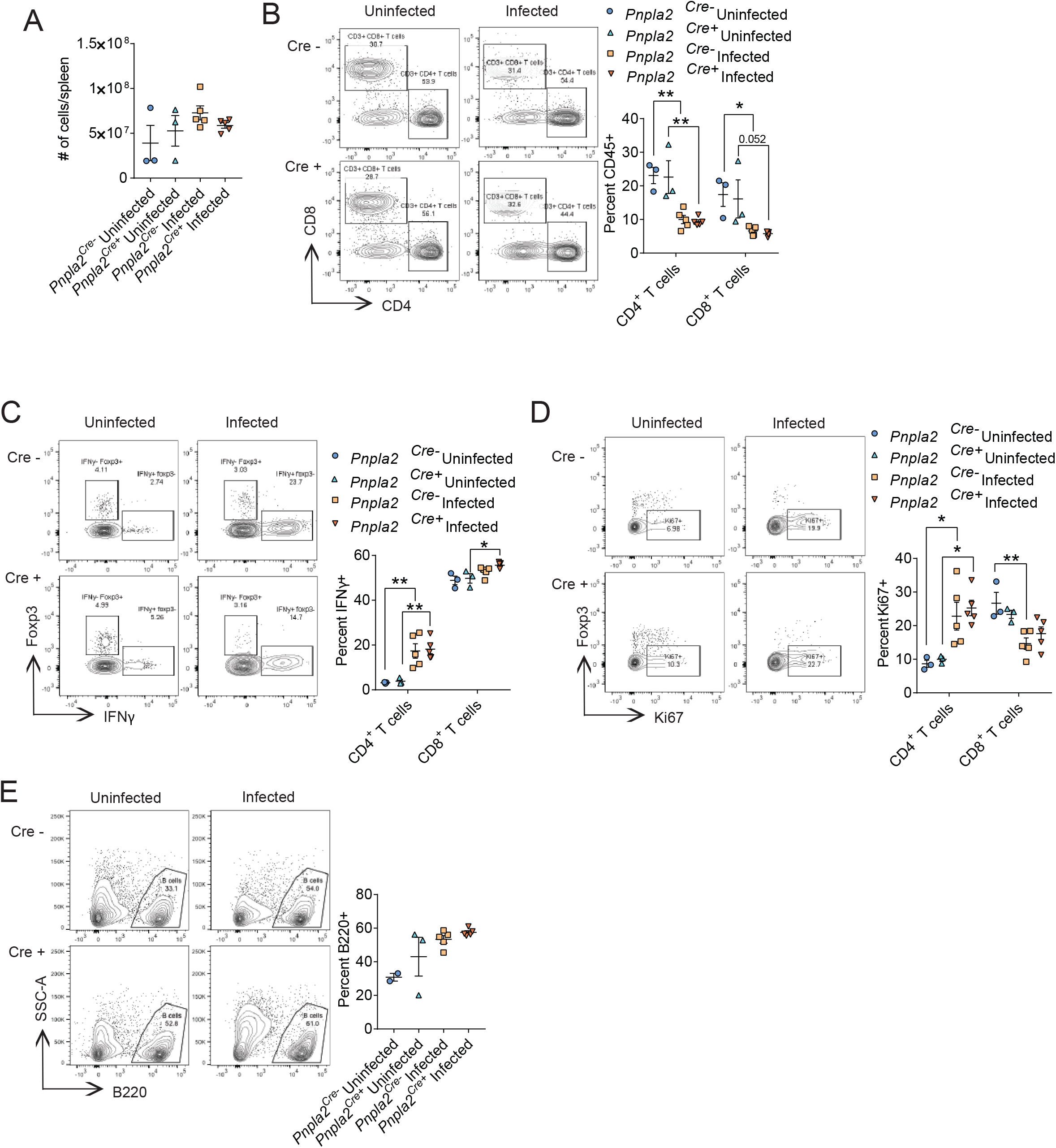
Adipose tissue wasting does not affect the spleen immune cell populations. Mice were intravenously infected with 500,000 *T. brucei*. Cells were isolated from gonadal white adipose tissue (GWAT) of uninfected *Pnpla2^flox/flox^;Fabp4^Cre+^* (*Pnpla2^Cre+^*) mice (n=3), uninfected *Pnpla2^flox/flox^;Fabp4^Cre-^* (*Pnpla2^Cre-^*) (n=2-3 depending on panel), infected *Pnpla2^Cre-^* (n=5) and infected *Pnpla2^Cre+^* (n=5) littermates at 6 days post-infection and immune cell populations were quantified. These are the same mice as in Fig. 3. (A) The total number of CD45^+^ cells isolated from spleens of each mouse. (B) Percent of CD45^+^ cells that are either CD4^+^ T cells or CD8^+^ T cells. (C) Percent of either CD4^+^ T cells or CD8^+^ T cells that are IFNγ^+^ (D) Percent of either CD4^+^ T cells or CD8^+^ T cells that are ki67^+^ (E) Percent of CD45^+^ cells that are B cells (B220^+^). * p<0.05, ** p<0.01, *** p<0.001, and *** p<0.0001 One-way ANOVA comparing the mean of each column to every other mean using Tukey correction for multiple comparisons was done on all graphs. Error bars represent +/- SEM. Frequencies displayed on gates are percent of parent gate which may be different than what is being displayed on graph which is listed above for each panel. Panels show one representative experiment.

**Supplementary Figure 6.**
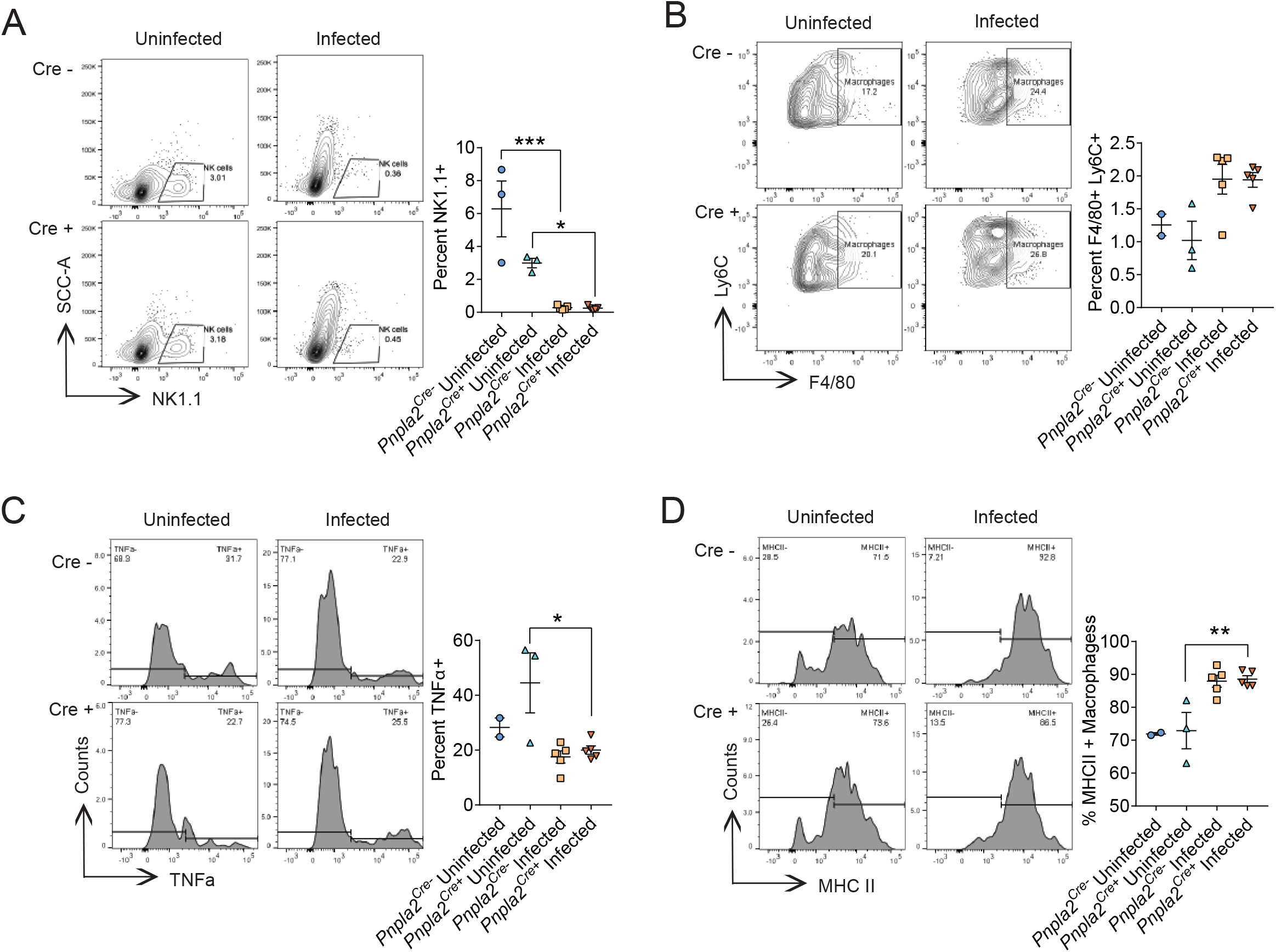
Innate immune cells of the spleen are not affected by adipose tissue wasting. Mice were intravenously infected with 500,000 *T. brucei*. Cells were isolated from gonadal white adipose tissue (GWAT) of uninfected *Pnpla2^flox/flox^;Fabp4^Cre+^* (*Pnpla2^Cre+^*) mice (n=3), uninfected *Pnpla2^flox/flox^;Fabp4^Cre-^* (*Pnpla2^Cre-^*) (n=2-3 depending on panel), infected *Pnpla2^Cre-^* (n=5) and infected *Pnpla2^Cre+^* (n=5) littermates at 6 days post-infection and immune cell populations were quantified. These are the same mice as in Fig. 3. (A) Percent of CD45^+^ cells that are Natural Killer cells (NK1.1^+^). (B) Percentage of CD45^+^ cells that are macrophages (CD11b^+^, Ly6C^int-high^, F4/80^+^, Ly6G^-^). (C) Percentage of macrophages that are TNFα^+^. (D) Percent of macrophages that are MHCII^+^. * p<0.05, ** p<0.01, *** p<0.001, and *** p<0.0001 One-way ANOVA comparing the mean of each column to every other mean using Tukey correction for multiple comparisons or Kruskal Wallis with a post Dunn test. Error bars represent +/- SEM. Frequencies displayed on gates are percent of parent gate which may be different than what is being displayed on graph which is listed above for each panel. Panels show one representative experiment.

**Supplemental Figure 7.**
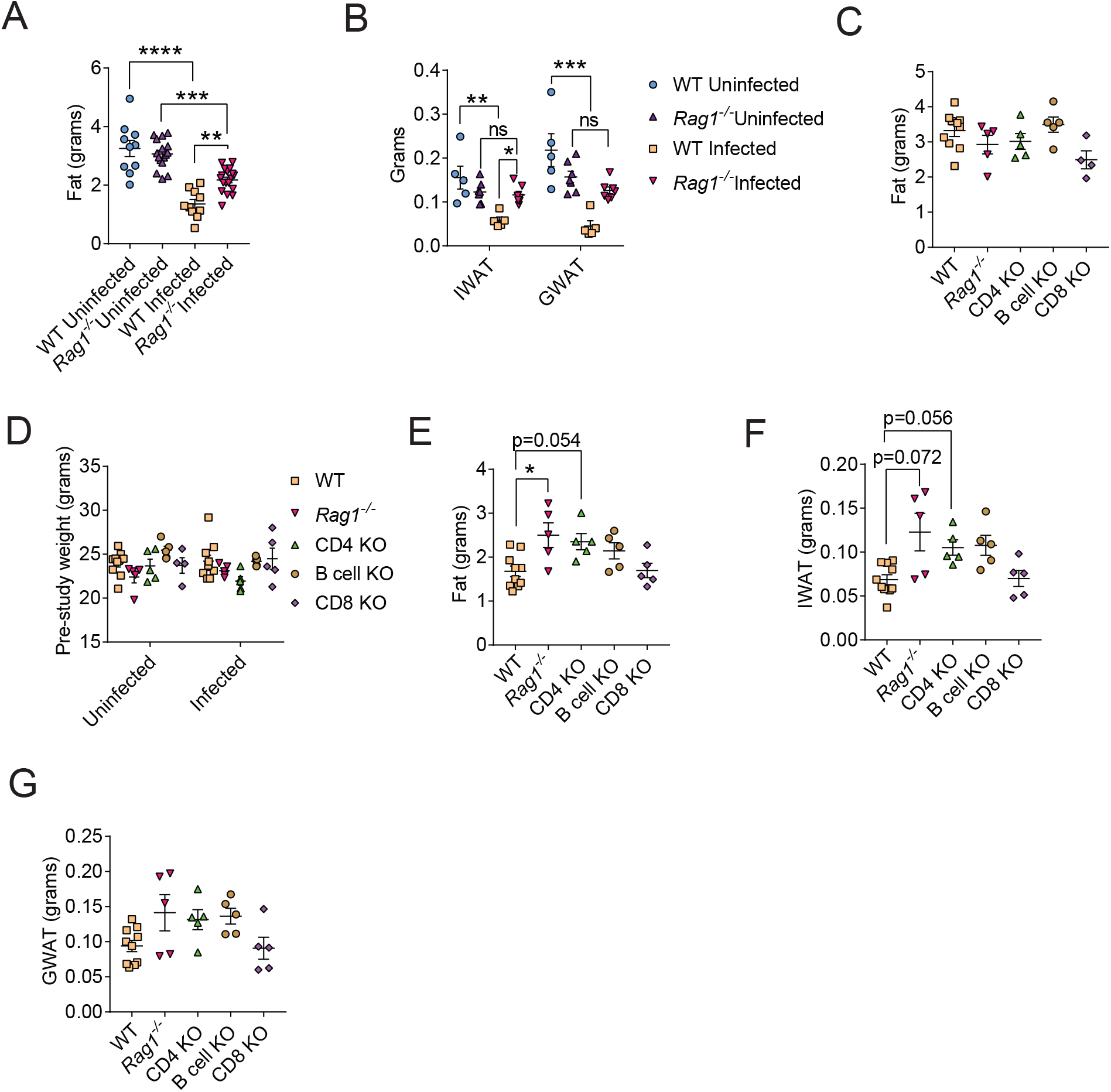
The adaptive immune system drives adipose tissue wasting. Mice were intravenously infected with 500,000 *T. brucei*. (A) Same mice displayed in Figure 4C showing MRI of fat mass from uninfected (n=10) and infected (n=10) C57BL/6J and uninfected (n=15) and infected (n=18) *Rag1* KO mice not normalized to pre-study weight. (B) Same mice displayed in Figure 4D showing IWAT and GWAT weights from uninfected (n=5) and infected (n=5) WT mice compared to uninfected (n=7) and infected (n=7) *Rag1* KO mice normalized to pre-study mouse weight. (C) MRI readings of fat tissue in grams of uninfected C57BL/6J (n=5) mice, B6.129S7-Rag1^tm1Mom^/J (n=5) mice (*Rag1* -/-), B6.129S2-Cd4^tm1Mak^/J (n=5) mice (CD4 KO), B6.129S2-Cd8a^tm1Mak^/J (n=5) mice (CD8 KO), and B6.129S2-Ighm^tm1Cgn^/J (n=5) mice (B cell KO). (D) Weights of infected mice from Figure 4E with their genotype weight matched uninfected controls at the time of infection. n=10 for both groups of WT mice and n=4-5 for all other groups. (E) MRI of fat tissue in grams of mice shown in Figure 4E with C57BL/6J (n=10) mice, B6.129S7-Rag1^tm1Mom^/J (n=5) mice (*Rag1* -/-), B6.129S2-Cd4^tm1Mak^/J (n=5) mice (CD4 KO), B6.129S2-Cd8a^tm1Mak^/J (n=5) mice (CD8 KO), and B6.129S2-Ighm^tm1Cgn^/J (n=5) mice (B cell KO) without normalizing to pre-study weight or uninfected mice. (F) Same mice as in the previous panel showing IWAT weights not normalized to pre-study weight of the mouse or the fat mass of uninfected mice. (G) Same mice ass in panel E of this figure showing GWAT weights not normalized to pre-study weight of the mouse for the fat mass of uninfected mice. * p<0.05, ** p<0.01, *** p<0.001, and *** p<0.0001. One-way ANOVA comparing the mean of each column to every other mean using Tukey correction for multiple comparisons or Kruskal Wallis with a post Dunn test. Error bars represent +/- SEM. Panel A shows two independent experiments combined, all other panels show one representative experiment.

**Supplemental Figure 8.**
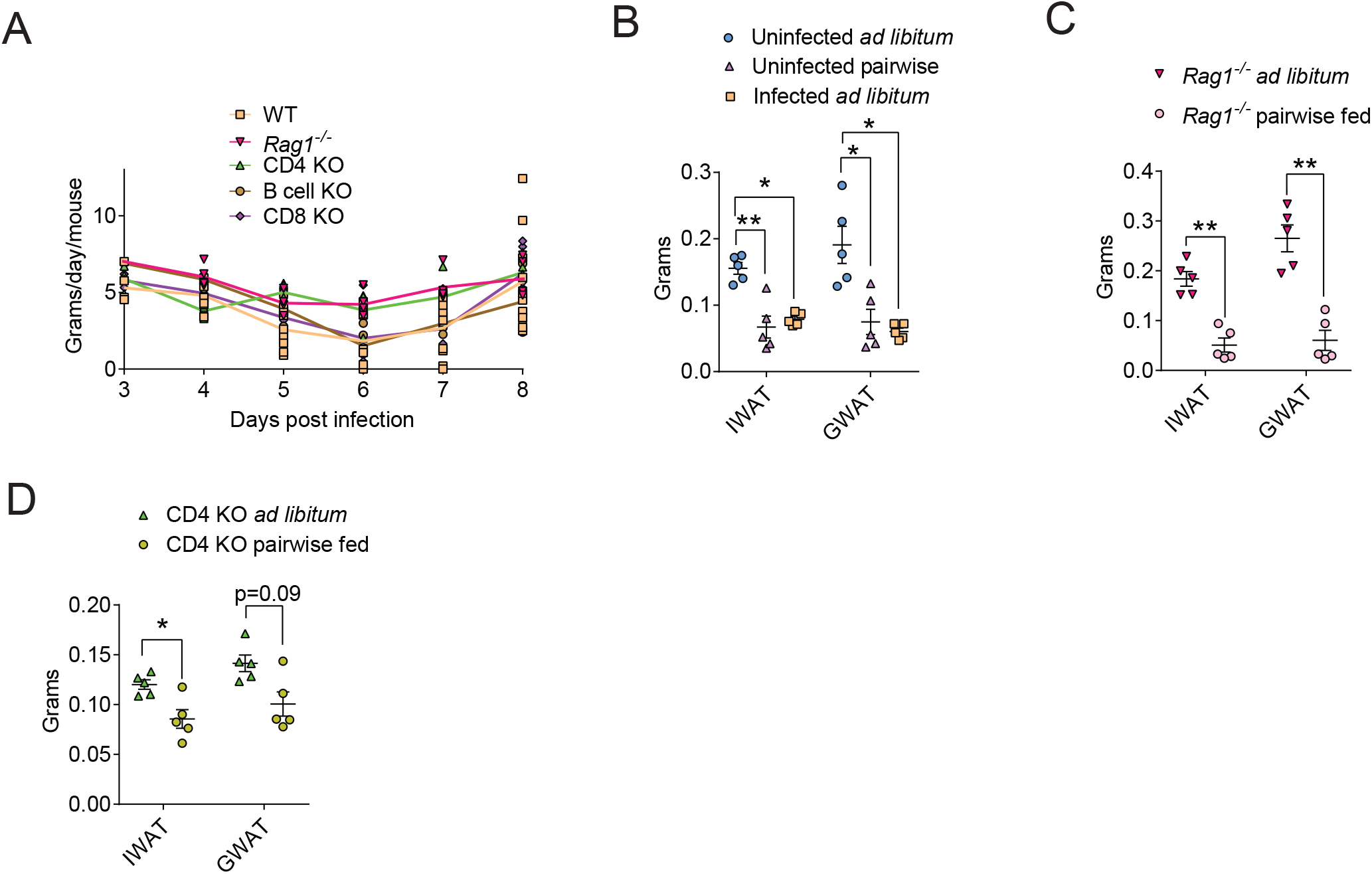
CD4^+^ T cells are necessary for the sickness induced anorexic response. Mice were intravenously infected with 500,000 *T. brucei* and the immune. (A) Same data as in Figure 5B showing individual data points for food intake of single-caged infected WT (n=5) mice, *Rag1*^-/-^ (n=5) mice, CD4 KO (n=5) mice, CD8 KO (n=5) mice, and B cell KO (n=5) mice. (B) Same data as in Figure 5E showing weights of both inguinal white adipose tissue (IWAT) and gonadal white adipose tissue (GWAT) from single caged, uninfected C57BL/6J fed ad libitum (n=5), uninfected C57BL/6J mice pairwise fed and given only the amount of food infected mice eat (n=5), and infected C57BL/6J fed ad libitum dissected at 7 days post-infection without normalizing to prestudy weight. (C) Fat pads without being normalized to starting weight of 12 week old *Rag1*^-/-^ mice that were infected and fed *ad libitum* (n=5) and infected pairwise fed mice (n=5) which were given the same amount of food as WT infected mice ate shown in Figure 5A. (D) Fat pads without being normalized to starting weight of 8 week old CD4 KO mice that were infected and fed *ad libitum* (n=5) and infected pairwise fed mice (n=5) which were given the same amount of food as WT infected mice ate shown in Figure 5A. (E) Titration curve of serum using ELISA to get a relative quantification of IgG antibodies from WT (n=5) mice, *Rag1*^-/-^ (n=4) mice, CD4 KO (n=5) mice, CD8 KO (n=5) mice, and B cell KO (n=5) mice. (F) Same mice as in panel E but quantifying IgM in the titration curve. * p<0.05, ** p<0.01, *** p<0.001, and *** p<0.0001. Kruskall Wallis tests were performed comparing the mean of each group to WT mice with Dunn test correcting for multiple comparisons for panel B. Mann-Whitney tests were also done on panels C and D. Error bars represent +/- SEM. Data depicts one representative experiment.

**Supplemental Figure 9.**
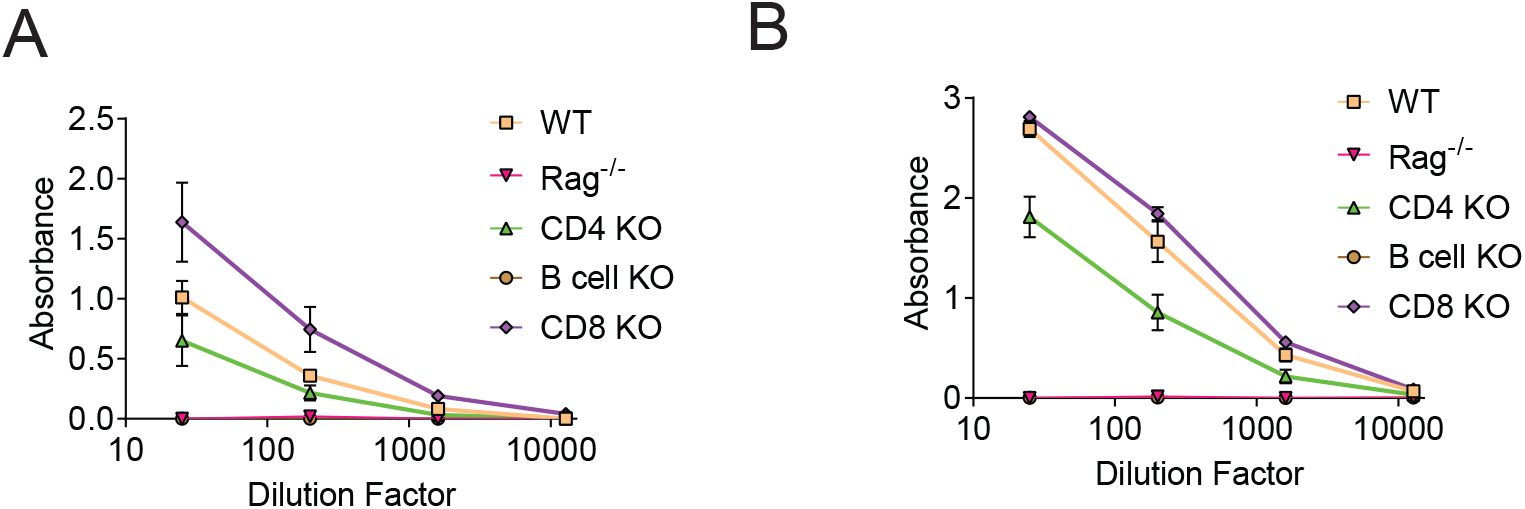
Decoupling resistance from adipose tissue wasting during *T. brucei* infection. Mice were intravenously infected with 500,000 *T. brucei* and the immune. (A) Titration curve of serum with ELISA to get a relative quantification of IgG antibodies from WT (n=5) mice, *Rag1* -/- (n=4) mice, CD4 KO (n=5) mice, CD8 KO (n=5) mice, and B cell KO (n=5) mice. (B) Same mice as in panel A but quantifying IgM. Error bars represent +/- SEM. Data depicts one representative experiment.

## Notes

### Competing Interest Statement

The authors have declared no competing interest.

